# The protective role of chloroplast NADH dehydrogenase-like complex (NDH) against PSI photoinhibition under chilling stress

**DOI:** 10.1101/2025.05.11.653304

**Authors:** Ko Takeuchi, Shintaro Harimoto, Yufen Che, Minoru Kumazawa, Hayato Satoh, Shu Maekawa, Chikahiro Miyake, Kentaro Ifuku

## Abstract

Chilling stress induces photosystem I (PSI) photoinhibition in various plants, severely impairing their growth. However, the mechanisms suppressing chilling-induced PSI photoinhibition remain unclear. This study aimed to identify factors preventing PSI photoinhibition by comparing two cucumber cultivars with different susceptibilities to PSI photoinhibition and chilling stress tolerance. In the chilling-sensitive cultivar, partial degradation of the CF_1_-γ subunit of chloroplast ATPase led to uncoupling of the thylakoid membrane. In addition, electron efflux from Fe-S clusters downstream of PSI was restricted under chilling stress, resulting in highly reduced Fe-S clusters. Notably, this PSI over-reduction in the chilling-sensitive cultivar was observed not only under chilling stress but also under fluctuating light conditions, limited CO_2_ conditions, and during the transition from darkness to light, suggesting that cyclic electron flow contributes to cultivar differences in PSI photoinhibition. Indeed, the chilling-tolerant cultivar exhibited higher activity of the chloroplast NADH dehydrogenase-like complex (NDH) and suppressed reactive oxygen species (ROS) accumulation during the early stages of chilling stress. In contrast, in the chilling-sensitive cultivar, destabilization of PSI–NDH supercomplex under chilling stress led to the loss of NDH activity, resulting in severe PSI over-reduction. This study provides evidence that NDH acts as a crucial electron sink to prevent PSI photoinhibition and provides new insights into the mechanisms underlying low-temperature stress tolerance.

## Introduction

Plants are exposed to a wide range of environmental stresses in nature. Photosystems are highly susceptible to environmental stress, leading to photoinhibition and reduced plant growth. Photosystem I (PSI) photoinhibition, caused by low temperatures or fluctuating light, severely affects plant growth because of the slow recovery of PSI (Kudoh and Sonoike, 2002; Sonoike, 2011; Zivcak *et al*., 2015; Li *et al*., 2018). Therefore, elucidating the mechanisms underlying PSI photoinhibition and the protective factors against PSI photoinhibition is crucial for enhancing plant stress tolerance and agricultural productivity.

PSI photoinhibition refers to PSI-specific damage independent of PSII photoinhibition. This phenomenon was first identified *in vivo* in cucumber, a typical chilling-sensitive plant. Under light conditions at temperatures below 10 °C, cucumber exhibits a decrease in photo-oxidizable PSI reaction-center chlorophyll P700 and PSI quantum yield, whereas PSII remains largely unaffected (Terashima *et al*., 1994). Subsequent studies confirmed that low-temperature (chilling stress)-induced PSI photoinhibition occurs in various plant species, particularly those adapted to warmer climates, indicating that this phenomenon is not exclusive to cucumbers (Havaux and Davaud, 1994; Erling Tjus *et al*., 1999; Kornyeyev *et al*., 2003; Govindachary *et al*., 2004; Kim *et al*., 2005; Scheller and Haldrup, 2005; Zhang Y. *et al*., 2024).

Chilling-induced PSI photoinhibition is driven by reactive oxygen species (ROS) because it does not occur under anaerobic or dark conditions (Terashima *et al*., 1998; Shimakawa *et al*., 2024). ROS-induced photo-oxidative damage degrades iron-sulfur (Fe-S) clusters (Sonoike *et al*., 1995; Sonoike, 1996*a*; Sonoike *et al*., 1997; Erling Tjus *et al*., 1998; Shimakawa *et al*., 2024) and PSI core proteins PsaA and PsaB (Sonoike and Terashima, 1994; Sonoike, 1996*b*; Sonoike *et al*., 1997; Ivanov *et al*., 1998; Erling Tjus *et al*., 1999; Nakano *et al*., 2010; Grebe *et al*., 2024). One contributing factor to chilling-induced PSI photoinhibition is that membrane destabilization and ROS production during chilling stress cause the uncoupling of thylakoids, preventing the suppression of electron influx into PSI (Peeler and Naylor, 1988; Terashima *et al*., 1991*a*,*b*). This uncoupling of thylakoids is associated with the release of the CF_1_ complex of chloroplast ATPase (Terashima *et al*., 1991*a*,*b*; Kono *et al*., 2022). However, the exact factors determining susceptibility to chilling-induced PSI photoinhibition have not been elucidated.

Recent studies have suggested that the most critical factor suppressing PSI photoinhibition under environmental stress is maintaining P700 and Fe-S clusters in an oxidized state (Miyake, 2020). If the Fe-S clusters (F_X_, F_A_, and F_B_) remain reduced for extended periods, the risk of ROS production increases (Furutani *et al*., 2023; Tiwari *et al*., 2024). In many angiosperms, thylakoid lumen acidification (ΔpH), which restricts electron influx into PSI by suppressing electron transport in Cyt *b*_6_*f*, is the most powerful P700 and Fe-S oxidation mechanism (Foyer *et al*., 1990; Anderson, 1992; Nishio and Whitmarsh, 1993; Hope, 2000; Baker *et al*., 2007; Kohzuma *et al*., 2009; Schöttler *et al*., 2015). This process is known as “photosynthetic control”, regulating the electron donation to PSI. In addition, reduction-induced suppression of electron flow (RISE) upstream of Cyt *b*_6_*f* as well as PSII photoinhibition can contribute to PSI oxidation (Shaku *et al*., 2016; Takeuchi *et al*., 2025). The oxidation of PSI was also achieved by activating the various reactions on the PSI acceptor side. Photorespiration serves as an important electron sink in C_3_ plants (Takagi *et al*., 2016; Wada *et al*., 2020). Electron flow from Fd to the plastoquinone (PQ) pool, called cyclic electron flow (CEF), also contributes to P700 oxidation (Shikanai, 2024). Furthermore, oxidized P700 (P700^+^) prevents Fe-S over-reduction via charge recombination (Trissl, 1997; Kou *et al*., 2015; Cherepanov *et al*., 2017; Degen and Johnson, 2024; Tiwari *et al*., 2024; Takeuchi et al., 2025).

Cucumber cultivars exhibit variation in their chilling stress tolerance (Lv *et al*., 2022; Takeuchi *et al*., 2022, 2024; Wang *et al*., 2022), providing an ideal model system to explore the physiological basis of PSI photoinhibition. Despite progress in identifying the multiple mechanisms that protect PSI, it remains unclear which mechanisms are decisive in conferring chilling tolerance. In this study, we aimed to dissect the physiological factors determining the extent of PSI photoinhibition and identify the key factors responsible for its suppression by comparing chilling-tolerant and chilling-sensitive cucumber cultivars.

## Materials and Methods

### Plant materials, growth conditions, and chilling stress treatment

Cucumber (*Cucumis sativus* L. cvs. ‘High Green 21’ [‘HG’] and ‘Homi 1-gou’ [‘HM’]) seeds were purchased from Saitama Genshu Ikuseikai (Saitama, Japan). Plants were grown in plastic pots with soil under a photoperiod of 16 h light (250–300 µmol photons m^−2^ s^−1^)/8 h darkness at 27 °C in a growth chamber. Fully expanded leaves from 2–3- week-old plants were used for all analyses. Cucumber plants were exposed to chilling stress at 4 °C in a cold room under light illumination of 180–250 µmol photons m^−2^ s^−1^. Photosynthetic parameters were measured after 1 h, 3 h, and 5 h of chilling stress. Plants not subjected to chilling treatment were used as control plants.

*Arabidopsis thaliana* (*A. thaliana*) wild type (ecotype Columbia-0) and the NADH dehydrogenase-like complex (NDH)-deficient mutant *pnsl1/ppl2* (Ishihara *et al*., 2007) were grown in soil under long-day conditions (16-h light/8-h dark) at 22 °C with 80–100 μmol photons m^−2^ s^−1^. Rosette leaves of 4– to 6-week-old plants were used for photosynthetic measurements.

### Measurement of chlorophyll content and ion leakage

Cucumber leaves were collected from the second or third fully expanded leaves (counted from the bottom) 2 days after 24 h of chilling stress. The samples were immediately frozen and ground into powder in liquid nitrogen. Total chlorophyll (Chl *a* + *b*) was extracted with 80% (v/v) acetone and was determined by spectrophotometry as previously described (Porra *et al*., 1989). Relative ion leakage was measured as previously described with minor modifications (Jambunathan, 2010). Leaves were collected 2 days after 24 h of chilling stress, and ion leakage was measured using COND LAQUAtwin (HORIBA, Japan). The ion leakage in the non-chilled control leaves of each cultivar was set to 1.

### Measurement of chlorophyll fluorescence, P700, and iron/sulfur clusters signals

Chl fluorescence and redox change of P700 were measured by induction curve or light curve methods using the Dual-PAM-100 measuring system (Heinz Walz GmbH, Effeltrich, Germany) at 23 °C as previously described (Takeuchi et al., 2022, 2024, 2025). The maximum quantum yield of PSII in the dark was calculated as F_v_/F_m_. The effective quantum yield of PSII was calculated as Y(II) = (F_m_′− F_s_)/F_m_′. The quantum yields of non-regulated and regulated energy dissipation in PSII were calculated as Y(NO) = F_s_/F_m_ and Y(NPQ) = F_s_/F_m_′− Y(NO), respectively. The F_o_ and F_m_ levels and the minimum and maximum fluorescence were measured after adaptation to the dark for at least 20 min. Light-chilled plants were adapted to darkness in a cold room (4 °C), whereas control plants were adapted at 23 °C. After the onset of actinic light (AL) illumination, saturation pulses (SPs) (300 ms and 20,000 µmol photons m^−2^ s^−1^) were applied to monitor F_m_′ (maximum fluorescence under light) and F_s_ (steady-state fluorescence under light). The redox changes in P700 were analyzed by monitoring the absorbance changes at 830 and 875 nm (Klughammer and Schreiber, 2008).

The ratio of P700^+^ under AL conditions was calculated as Y(ND) = P/P_m_. The fraction of P700 that could not be oxidized by applying SPs to the overall P700 was calculated as Y(NA) = (P_m_ − P_m_′)/P_m_. Y(I) was calculated as (P_m_′ − P)/P_m_ (Klughammer and Schreiber, 1994). P_m_ was determined by applying SPs after far-red (FR) light illumination. P_m_′ represents the maximum level of P700^+^ under AL. P represents the steady-state P700^+^ level recorded immediately before the application of SPs. Notably, Y(I) and Y(NA) depend only on the step of the P700 redox cycle that is rate-limited during the application of SPs and do not reflect the quantum yield of PSI under AL (Furutani et al., 2022).

To measure Chl fluorescence and the redox state of P700 under low CO_2_ conditions, light curve methods were adapted under conditions wherein air was vented into soda lime for CO_2_ absorption, and CO_2_ concentrations were monitored using the GM70 CO_2_ sensor (VAISALA, Finland) and maintained below 30 ppm.

The redox states of Fe–S clusters, including F_X_, F_A_/F_B_, and Fd, were measured at 23 °C using the Dual-KLAS-NIR spectrophotometer (Heinz Walz GmbH, Germany) as described previously (Klughammer and Schreiber, 2016; Schreiber, 2017). Maximum Fe–S signals and reduced Fe–S levels under AL (300 µmol photons m^−2^ s^−1^) before and after exposure to the chilling stress were measured as previously described (Ohnishi et al., 2023; Maekawa et al., 2024).

### Measurement of NDH activity

NDH activity was assessed by monitoring the transient increase in Chl fluorescence after AL was turned off using MINI-PAM (Heinz Walz GmbH, Effeltrich, Germany). Leaves were dark-adapted for at least 20 min before exposure to AL (60 μmol photons m^−2^ s^−1^) for 4–5 min. The subsequent transient increase in Chl fluorescence was monitored under the measuring light immediately after the AL was turned off.

### Electrochromic shift (ECS) measurements

ECS was measured using Dual-PAM 100 equipped with a P515/535 module (Heinz Walz GmbH, Effeltrich, Germany). Two- to three-week-old cucumbers were exposed to 1 h or 3 h of chilling stress. After 20 min of dark adaptation at 4 °C (control plants at 23 °C), proton motive force (*pmf*) was determined by ECS under ambient conditions. ECS was recorded by exposing the samples to a minute of AL at 325, 600, and 1200 µmol photons m⁻² s⁻¹ in this order, with dark relaxation periods between each AL intensity to measure ECSt. ECSt was normalized to the maximum change in absorbance at 515 nm observed in control plants.

### Photorespiration activity

Photorespiration activity was measured as previously described (Sejima *et al*., 2016; Hanawa *et al*., 2017). O_2_ exchange and Chl fluorescence were simultaneously monitored. Leaf discs were placed in an O_2_-electrode chamber (Hansatech, UK), and Chl fluorescence was monitored using Junior-PAM (Heinz Walz GmbH, Effeltrich, Germany) through a light-guided plastic fiber inserted into the O_2_ electrode. The temperature was maintained at 25 °C for the control measurement and at 15 °C for the measurement after chilling stress, using a circulating temperature-controlled water jacket. Light (1200 μmol photons m^−2^ s^−1^) was illuminated from the top of the O_2_-electrode chamber. The post-illumination transient O_2_-uptake rate reflected the activities of ribulose 1,5-bisphosphate oxygenase (Hanawa *et al*., 2017).

### Isolation of thylakoid membranes

To isolate thylakoid membranes, fresh cucumber leaves (5 g) were cut into small stripes and ground in 50 mL ice-cold buffer containing 0.3 M sorbitol, 10 mM NaCl, 5 mM MgCl_2_, 0.1% (w/v) bovine serum albumin, 2 mM Na-ascorbate, 10 mM EDTA-Na_2_, protease inhibitor cocktail (Protease Inhibitor Cocktail Set I, Animal-derived-free, FUJIFILM, Japan), and 40 mM HEPES-KOH (pH 7.5) using a polytron homogenizer (PT 10-35GT, Kinematica company, Switzerland) at a line voltage of 6 for 10 s. The homogenate was filtered through a single layer of Miracloth (EMD Millipore, U.S.A.) and centrifuged at 2000×g for 2 min at 4 ℃. The pellet was resuspended in a medium containing 0.33 M sorbitol, 15 mM NaCl, 7.5 mM MgCl_2_, and 40 mM HEPES-KOH (pH 7.5).

### Clear-native PAGE (CN-PAGE), blue-native PAGE (BN-PAGE), SDS-PAGE, and immunoblot analysis

CN-PAGE was performed as described previously (Che *et al*., 2020). After solubilizing the thylakoid membranes with *n*-dodecyl-*β*-D-maltoside (β-DM) on ice for 10 min, the insoluble material was removed by centrifugation at 14,500 rpm for 10 min. Thylakoid membrane samples containing equal amounts of Chl were loaded onto a native PAGE 4– 16% Bis-Tris gel (Invitrogen, U.S.A.). Electrophoresis was performed at 100 V at 4°C. For 2D-CN/SDS-PAGE analysis, CN-PAGE gel lanes were excised and soaked in SDS sample buffer containing 2.5% (v/v) β-mercaptoethanol for 15 min at 23 °C and 15 min at 65 °C. Each lane with denatured proteins was placed on the top of 1.5-mm-thick 12% or 15% wide-range gels and electrophoresed at 100 V. The gels were subjected to immunoblotting. Proteins comparable to 10 μg Chl were loaded for CN-PAGE analysis.

BN-PAGE was performed as described by (Järvi *et al*., 2011). After solubilizing the thylakoid membranes with β-DM on ice for 1 min, the insoluble material was removed by centrifugation at 15,000 rpm for 2 min. Thylakoid membrane samples were loaded onto a Native PAGE 4–13% gradient gel. Electrophoresis was performed at 40 V at 4 °C. For 2D-BN/SDS-PAGE analysis, BN-PAGE gel lanes were excised and soaked in 1% SDS solubilization buffer containing 0.05 M dithiothreitol for 30 min at 23 °C. Each lane with denatured proteins was placed on the top of 1.0-mm-thick 12% wide-range gels and electrophoresed at 50 mA. The gels were subjected to immunoblotting. Proteins comparable to 5 μg Chl were loaded for BN-PAGE analysis.

SDS-PAGE and immunoblot analyses were performed as described previously (Che *et al*., 2020). Thylakoid membrane protein samples were separated on 12% or 15% wide-range gels and transferred to 0.22 or 0.45 μm PVDF membranes. The membranes were blocked with 5% (w/v) skim milk and subsequently incubated with specific antibodies against the indicated proteins. Detection was performed using Amersham ECL Prime Western Blotting Detection Reagent (GE Healthcare). Anti-NdhH, PnsL1, NdhT, PsaA, PsaB, Atpβ (CF1-β), Atpγ (CF1-γ), and Cyt *f* specific antibodies were purchased from Agrisera.

### Measurement of O_2_^•–^ content

O_2_^•–^ content was determined as described previously (Tachibana *et al*., 2024; Takeuchi *et al*., 2025). Fresh leaves were harvested before and immediately after the chilling stress treatment. The liquid nitrogen-frozen samples were homogenized and mixed well with 1.5 mL phosphate buffer (pH 7.8) containing 10 μM hydroxylammonium chloride and 100 μM EDTA-Na_2_. The homogenate was centrifuged at 5,000 rpm for 5 min at 4 °C, and 500 μL of supernatant was transferred into fresh tubes. Subsequently, 1 mL of 17 mM sulfanilamide (in 30% (v/v) acetic acid) and 1 mL of 7 mM naphthalene diamine dihydrochloride (in distilled water) were added into each tube in that order, and the solution mix was incubated for 10 min at 37 °C. Next, 3 mL of diethyl ether was added to each tube, and the resultant solution was centrifuged at 5,000 rpm for 5 min at 23 °C. The absorbance of each sample was recorded at 540 nm. Calibration curves were established from 0–4 μM NO_2_^−^. The coefficient of determination *r*^2^ between absorbance at 540 nm and [NO_2_^−^] was 0.999. Based on the reaction 2O_2_^•–^ + H^+^ + NH_2_OH → H_2_O_2_ + H_2_O + NO_2_^−^, the concentration of O_2_^•–^ was calculated from the calibration curve according to [O_2_^•–^] = 2[NO_2_^−^] (μM).

### RNA sequencing

RNA was extracted from the second fully expanded leaf (counted from the bottom) before and immediately after 1 or 3 h of chilling stress, as previously described (Che *et al*., 2020). Raw paired-end RNA-seq reads obtained using the DNBSEQ-G400 sequencer (MGI Tech Co., Ltd.) were trimmed with the Trimmomatic software (v0.39, http://www.usadellab.org/cms/?page=trimmomatic) with the forward adapter sequence 5’-AAGTCGGAGGCCAAGCGGTCTTAGGAAGACAA-3’ and the reverse adapter sequence 5’-AAGTCGGATCGTAGCCATGTCGTTCTGTGAGCCAAGGAGTTG-3’.

Libraries were prepared as described earlier (Kılıç *et al*., 2023). The quality of the trimmed reads was checked using FastQC (v0.11.8, https://www.bioinformatics.babraham.ac.uk/projects/fastqc/). The cleaned reads were mapped to the reference genome (Chinese Long v3) using Hisat2. Heatmaps were created using Morpheus software (https://software.broadinstitute.org/morpheus/).

### Statistical analysis

Statistical analyses were based on Tukey-Kramer’s multiple comparison test after one-way ANOVA or Student’s *t*-test. All calculations were performed using at least three independent biological replicates.

## Results

### Cultivar differences in chilling-induced PSI photoinhibition in cucumbers

To confirm the relationship between chilling stress tolerance and PSI photoinhibition, we compared the physiological responses to chilling stress in two cucumber cultivars: chilling-sensitive ‘HG’ and chilling-tolerant ‘HM’ (Takeuchi *et al*., 2022). HG exhibited severe leaf chlorosis, a significant reduction in chlorophyll content, and increased electrolyte leakage compared with that in HM after chilling stress (Fig. 1A). Fv/Fm declined slightly in both cultivars from approximately 0.8 to 0.6 within the first 5 h of chilling stress, with no marked difference between the two cultivars (Fig. 1B). This suggests a similar level of PSII photoinhibition. In contrast, PSI photoinhibition, as determined by Pm (the maximum oxidizable P700), progressed more rapidly in the chilling-sensitive cultivar HG than in the chilling-tolerant cultivar HM. Following chilling stress, photo-oxidizable P700 decreased to approximately 30% in HG, while HM retained 80% of active P700. Active Fe-S clusters also decreased significantly in the chilling-sensitive cultivar HG after chilling stress, whereas Fe–S clusters did not decrease significantly in HM (Fig. 1B). These results indicate that PSI photoinhibition progressed rapidly in the HG, leading to severe growth inhibition. In contrast, PSI photoinhibition was suppressed in the HM, correlating with its higher chilling tolerance.

**Figure 1.**
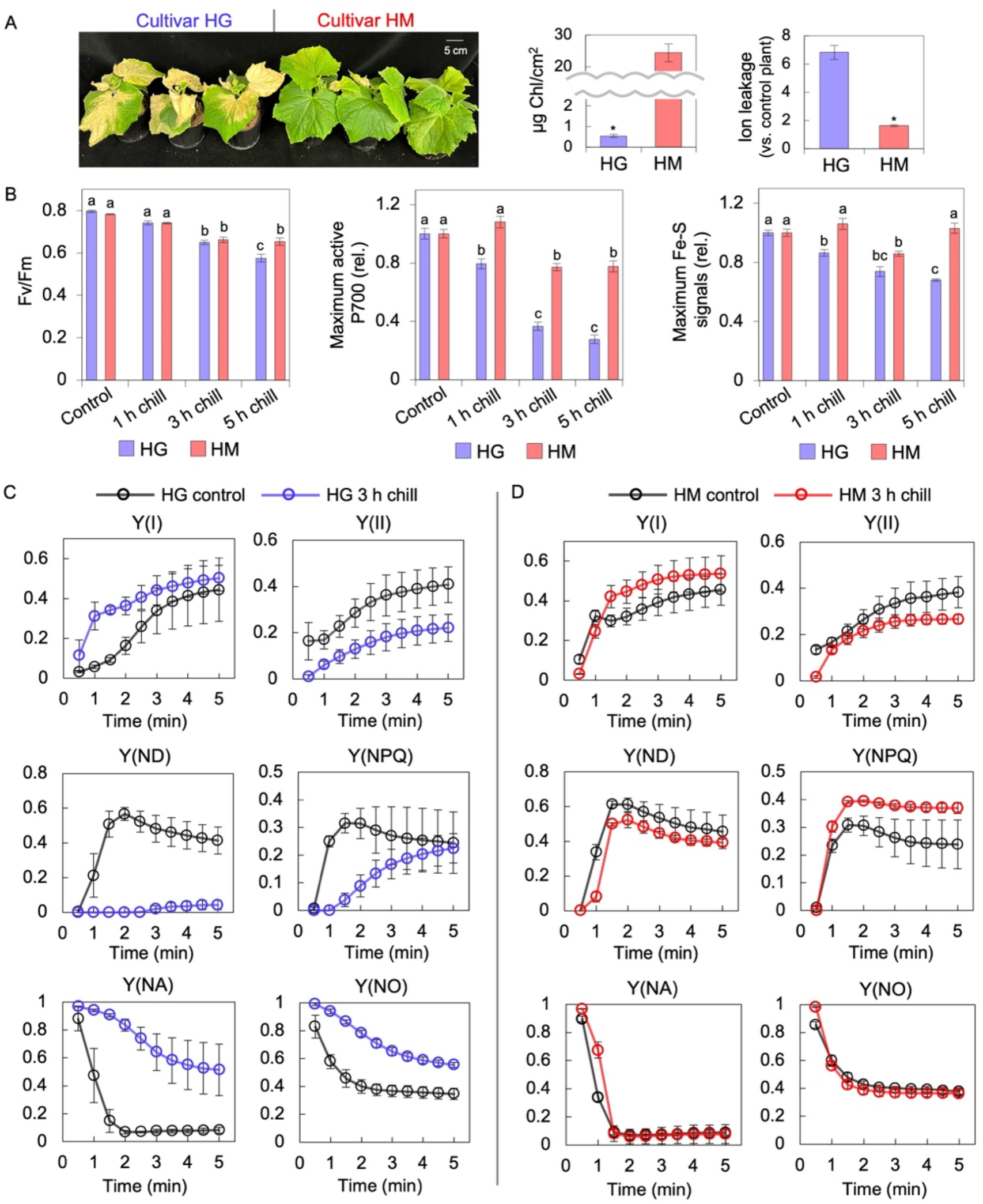
Phenotypes and photoinhibition caused by chilling stress in two cucumber cultivars. (A) Cucumbers (cvs. ‘HG’ and ‘HM’) were subjected to chilling stress at 4 °C under light conditions for 24 h. After chilling stress, plants were returned for 2 days to their original chamber at 27 °C to assess their recovery. Subsequently, total Chl content and ion leakage were measured. Values are the mean ± SE, n = 3–4, biological replicates. Asterisks denote statistical significance (*p*<0.01, Student’s *t*-test). (B) The maximum quantum yield of PSII (Fv/Fm), maximum amount of photo-oxidizable P700 (Pm), and maximum Fe-S cluster signals were measured spectroscopically using Dual-KLAS-NIR before and after 1 h, 3 h, and 5 h of chilling stress. Plants were dark-adapted for at least 20 min before measurements. Pm and Fe-S cluster signals were normalized to the values before chilling stress for each cultivar. Values are the mean ± SE, n = 3–6, biological replicates. Different letters indicate statistically significant differences (*p*<0.05, Tukey–Kramer’s multiple comparison tests after one-way ANOVA). (C, D) Chl fluorescence and P700 redox states were analyzed before and after 3 h of chilling stress. Chilling-treated plants were dark-adapted for at least 20 min at 4 °C, whereas control plants were adapted at 23 °C. Chl fluorescence and Chl absorption under the induction phase of photosynthesis were measured using Dual-PAM 100 at 23 °C under the illumination of AL (325 µmol photons m^−2^ s^−1^) for 5 min. Values are the mean ± SE, n = 3, biological replicates.

To further assess PSI photoinhibition, we measured the photosynthetic activity after 3 h of chilling stress (Fig. 1C). Y(ND), the fraction of oxidized P700 (P700⁺), nearly disappeared in the chilling-sensitive cultivar HG, whereas Y(NA), reflecting acceptor-side limitations in PSI during saturation pulses (SP), significantly increased after chilling stress (Fig. 1C). HG also showed a decline in Y(II) and the induction of Y(NPQ), indicating reduced PSII electron transport and non-photochemical quenching. In contrast, the chilling-tolerant cultivar HM maintained Y(ND) and Y(NA) at control levels even after chilling stress (Fig. 2B). HM also showed a higher Y(NPQ) than that of HG (Fig. 1D), suggesting a higher proton motive force (*pmf*).

**Figure 2.**
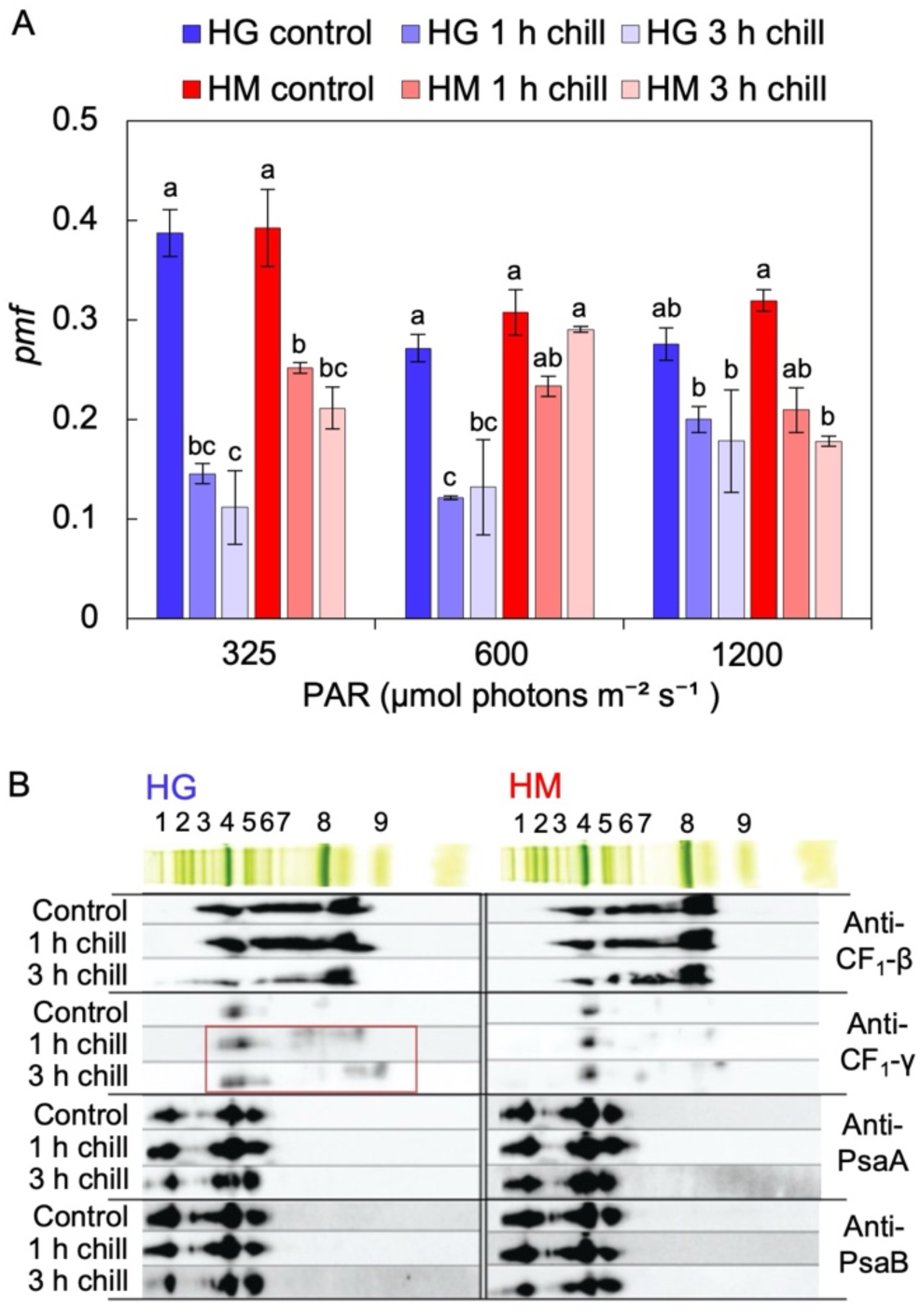
Impact of chilling stress on the *pmf* and stability of proteins in thylakoids. (A) Effect of chilling stress on *pmf*. Two- to three-week-old cucumbers (chilling-sensitive cv. ‘HG’ and chilling-tolerant cv. ‘HM’) were exposed to 1 h or 3 h of chilling stress. After 20 min of dark adaptation at 4 °C (control plants at 23 °C), *pmf* was determined by electrochromic shift (ECS) under ambient conditions using Dual-PAM 100 equipped with a P515/535 module. ECS was recorded by exposing the samples to a minute of AL at 325, 600, and 1,200 µmol photons m⁻² s⁻¹ in that order, with dark relaxation periods between each AL intensity to measure ECSt. ECSt was normalized to the maximum change in 515 nm absorbance observed in the control plants. Values are the mean ± SE, n = 3, biological replicates. Different letters indicate statistically significant differences (*p*<0.05, Tukey–Kramer’s multiple comparison tests after one-way ANOVA against each light intensity). (B) Assembly status of the CF_1_ subunit of chloroplast ATPase complexes and PSI core subunits (PsaA and PsaB). Two- to three-week-old cucumbers (chilling-sensitive cv. ‘HG’ and chilling-tolerant cv. ‘HM’) were treated at 4 °C for 1 h or 3 h. Isolated thylakoids separated by CN-PAGE before and after chilling stress (Supplementary Fig. S1) were analyzed by 2D-SDS-PAGE and subjected to immunoblotting with specific antibodies (anti-CF_1_-β, -CF_1_-γ, -PsaA, and -PsaB). The CN-PAGE images shown as an example above the immunoblotting data represent those obtained after 3 h of chilling treatment in each cultivar. The red outline in the figure represents the signals of the CF_1_-γ subunit that have been destabilized by the chilling stress. 1: PSI–NDH supercomplex; 2: PSII–LHCII supercomplex; 3: PSI + LHCI; 4: PSI– LHCI, PSII core dimer, and ATPase; 5: PSI core monomer; 6: PSII core monomer and Cyt *b*_6_*f*; 7: LHCII assembly; 8: LHCII trimer; and 9: LHCII monomer.

### Uncoupling of thylakoids induced by chilling stress

Previous studies have indicated that chilling stress induces the uncoupling of thylakoid membranes associated with the dissociation of the CF_1_ complex of the chloroplast ATPase, thereby preventing P700 oxidation (Y(ND)) (Peeler and Naylor, 1988; Terashima *et al*., 1991*a*,*b*). To evaluate the coupling state of cucumber thylakoids, we measured the *pmf* in chilled leaves at 23 °C using the P515 signal, which reflects ECS in absorption by pigments (chlorophylls and carotenoids). *Pmf* was measured by exposing the dark-adapted leaves to a minute of AL at 325, 600, and 1,200 µmol photons m⁻² s⁻¹ in that order, with dark intervals between each AL to measure ECSt. After the chilling stress, *pmf* declined significantly in both cultivars at all AL intensities (Fig. 2A). However, the chilling-tolerant cultivar HM tends to have a higher *pmf* than the chilling-sensitive cultivar HG at 325 and 600 µmol photons m⁻² s⁻¹. At 1200 µmol photons m⁻² s⁻¹, the difference in *pmf* between cultivars was reduced, likely because of temperature acclimation during measurement and light-induced warming. To test the stability of the CF_1_ and PSI core proteins in the thylakoid membrane, thylakoids separated by CN-PAGE before and after chilling stress (Supplementary Fig. S1) were analyzed by 2D-SDS-PAGE (Fig. 2B). The CF_1_-γ signal in the chilling-sensitive cultivar HG was widely detected at a lower molecular weight position than its original band 4 position in the thylakoids after 1 h and 3 h of chilling stress. This suggests that the stability of CF_1_-γ in the thylakoid membrane was disrupted by chilling stress, as previously reported (Terashima *et al*., 1991*a*,*b*). The observed destabilization of CF_1_-γ in HG would cause the differences in *pmf* and Y(NPQ) between the cultivars (Fig. 1C and 2A) (Takagi *et al*., 2017). In contrast, the stabilities of CF_1_-β, PsaA, and PsaB were reduced after 3 h of chilling stress in both cultivars (Fig. 3B). These results suggested that although the chilling-tolerant cultivar HM maintained CF_1_ stability to a certain extent compared with that in HG, photosynthetic control, a key factor in regulating P700 oxidation on the PSI donor side, became less functional in both cultivars under chilling stress.

**Figure 3.**
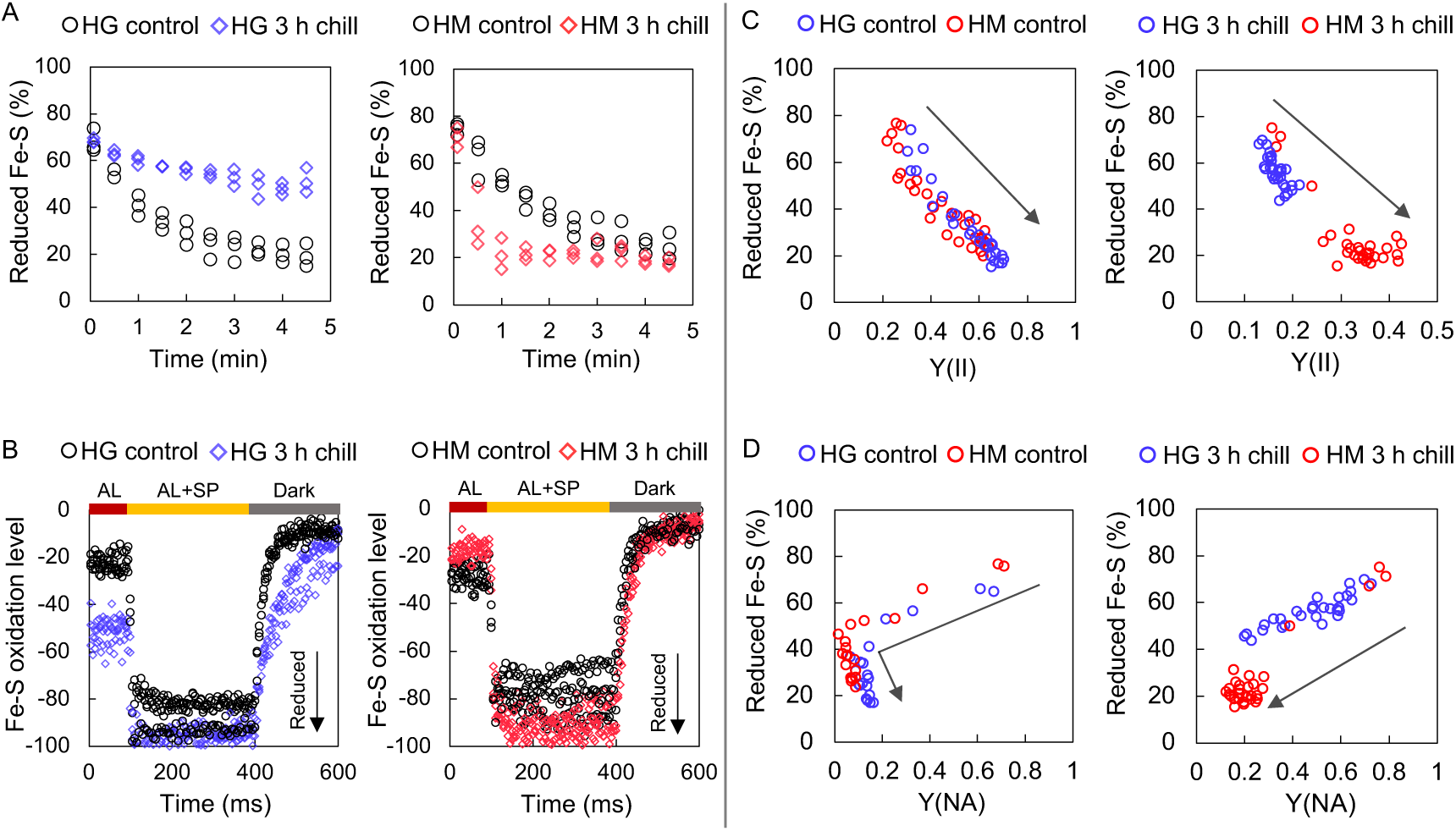
Reduction levels of Fe-S clusters under chilling stress. Two- to three-week-old cucumbers (chilling-sensitive cv. ‘HG’ and chilling-tolerant cv. ‘HM’) were treated at 4 °C for 3 h and were adapted to darkness for at least 20 min in a cold room (4 °C). Plants without chilling stress were used as controls. (A) Reduced levels of Fe-S clusters under the induction phase (dark-to-light transitions) of photosynthesis were analyzed spectroscopically using Dual-KLAS-NIR. AL (300 µmol photons m^−2^ s^−1^) was applied under ambient conditions (23 °C) for 5 min. (B) Re-oxidation of Fe-S clusters was assessed after 5 min of AL (300 µmol photons m^−2^ s^−1^) illumination by applying SP to reduce Fe-S clusters, followed by monitoring re-oxidation kinetics in the dark (400–600 ms) using Dual-KLAS-NIR. n = 3, biological replicates. (C) Reduced Fe-S levels under the 5-minute induction phase (dark-to-light transitions) of photosynthesis (Fig. 3A) were plotted against Y(II) measured simultaneously. The left graph shows data from control plants (without the chilling stress treatment), and the right graph shows data from chilling-treated plants (subjected to 3 h of chilling). The arrows in the graph represent the time course of the 5-min analysis. n = 3, biological replicates. (D) Reduced Fe-S levels (Fig. 3A) were plotted against Y(NA) measured simultaneously. The left graph shows data from control plants (without the chilling stress treatment), and the right graph shows data from chilled plants (3 h of chill). The arrows in the graph represent the time course of the 5-minute analysis. n = 3, biological replicates.

### Over-reduction of Fe-S clusters in the chilling-sensitive cultivar

To investigate additional factors influencing PSI photoinhibition on the PSI acceptor side, we measured the redox state of Fe-S clusters using Dual-KLAS-NIR. Under normal conditions, when light begins to irradiate dark-adapted leaves, the reaction that consumes the electrons of Fd becomes activated, leading to a gradual decrease in reduced Fe-S clusters (Fig. 3A, control) (Kadota *et al*., 2019). However, in the chilling-sensitive cultivar HG, more than 45% of Fe-S clusters remained reduced even 5 min after the onset of AL illumination after chilling stress, indicating persistent over-reduction of Fe-S. In contrast, in the chilling-tolerant cultivar HM, Fe-S clusters were rapidly oxidized even after chilling stress, and the oxidation of Fe–S clusters was faster than in the control (Fig. 3A). The over-reduction of Fe-S clusters after chilling stress in HG was further assessed by measuring the re-oxidation rate of the reduced Fe-S in the dark after turning off the light. The re-oxidation rate of reduced Fe-S in the dark reflects the activation level of the electron efflux reactions downstream of Fd (Schreiber, 2017). After 3 h of chilling stress, the chilling-sensitive cultivar HG exhibited delayed Fe-S re-oxidation compared with that in the control, indicating restricted electron efflux from Fd, whereas the chilling-tolerant cultivar HM showed no delay (Fig. 3B).

The reduced levels of Fe-S, measured every 30 s for 5 min after AL illumination, were plotted against Y(II), which was measured simultaneously (Fig. 3C). Under control conditions, a clear inverse correlation was evident with no cultivar differences (Fig. 3C, left). Immediately after light exposure, Fe-S was in a reduced state, and Y(II) was low. As time progressed, Fe-S was oxidized, and Y(II) increased. This indicates that during the induction phase of photosynthesis, Y(II) fully follows the redox state of the PSI acceptor side. After 3 h of chilling stress, the chilling-sensitive cultivar HG exhibited high levels of reduced Fe-S with extremely low Y(II) (0.1–0.2) (Fig. 3C, right), whereas the chilling-tolerant cultivar HM tended to keep Fe-S remaining oxidized with Y(II) around 0.3–0.4 (Fig. 3C, left). In Fig. 3D, the reduction levels of Fe-S were plotted against Y(NA). This is important because Y(NA) is often used as an indicator of the limitation of the acceptor side of P700 under AL; however, Y(NA) depends only on the rate-limiting step of the P700 redox cycle under SP, and it is unclear whether it accurately represents the over-reduction of PSI under AL (Furutani et al., 2022). Basically, reduced Fe-S levels correlated positively with Y(NA), indicating that PSI over-reduction was reflected in Y(NA) (Fig. 3D) (Maekawa *et al*., 2024). Specifically, under control conditions, during the initial few minutes after the onset of AL, Fe-S became oxidized, and Y(NA) decreased accordingly, dropping temporarily to nearly zero when approximately 40% of the Fe-S clusters were in a reduced state (Fig. 3D, left). This was likely because of the suppression of *pmf*-mediated electron influx into P700 before the activation of the electron efflux pathway after Fd, as indicated by transient increases in Y(ND) and Y(NPQ) at 2 minutes after AL illumination (Fig. 1C, D, control). By 5 minutes after AL illumination, with the activation of the electron transport pathway beyond FNR, both the Fe-S reduction levels and Y(NA) stabilized to similar values (Fig. 3D, left). After 3 h of chilling stress, the chilling-sensitive cultivar HG showed higher levels of reduced Fe-S and Y(NA) than those in HM (Fig. 3D, right). These results demonstrated that restricted electron efflux from Fd under chilling stress led to the over-reduction of Fe-S and a decline in Y(II) in the chilling-sensitive cultivar HG; however, the chilling-tolerant cultivar HM maintained an oxidized PSI acceptor side even under chilling stress through the oxidation of Fe-S (Fig. 3).

### Difference in Alternative electron flow (AEF) activity determines chilling stress tolerance

As chilling stress inhibits the Calvin–Benson–Bassham (CBB) cycle, alternative electron flow (AEF) becomes the primary electron sink for the oxidation of Fe-S (Kingston-Smith *et al*., 1997; Zhang Y. *et al*., 2024). To assess the differences in AEF activity, we restricted the CO_2_ supply to downregulate the CO_2_ assimilation pathway of the CBB cycle and monitored P700 redox states at normal temperatures. Notably, the chilling-sensitive cultivar HG exhibited lower Y(ND) and higher Y(NA) than those in HM, indicating a greater over-reduction in PSI under low CO_2_ conditions (Fig. 4). Importantly, no cultivar differences were observed in PSII parameters, such as Y(II), Y(NPQ), and Y(NO) (Fig. 4). These results suggested that when CO_2_ assimilation is limited, the chilling-sensitive cultivar HG is more susceptible to PSI over-reduction because of insufficient AEF activity downstream of Fe-S.

**Figure 4.**
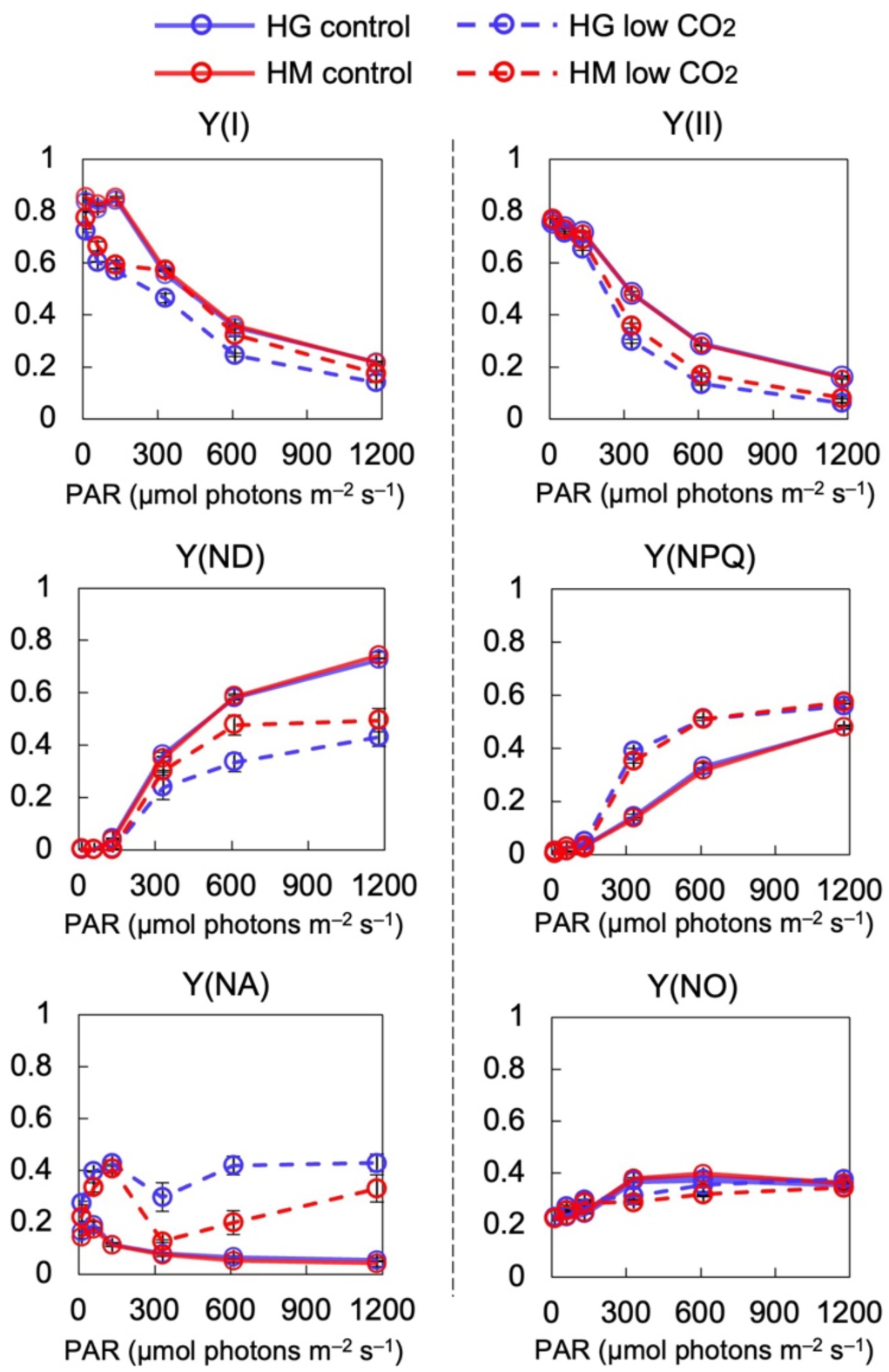
Impact of CO_2_ limitation on P700 oxidation in cucumber. Chl fluorescence and P700 redox states were measured under various light intensities in ambient (control) or low CO_2_ (< 30 ppm) conditions at 23 °C using Dual-PAM 100. The intact leaves of 2–3-week-old cucumbers (chilling-sensitive cv. ‘HG’ and chilling-tolerant cv. ‘HM’) were analyzed. Values are the mean ± SE, n = 3–4, biological replicates.

To examine the differences in AEF at the gene expression level between cultivars, we analyzed the gene expression changes in all nuclear-encoded CEF-related and photorespiration-related genes before and after chilling stress using RNA-seq. Heatmap analysis revealed that the expression of these genes tended to increase in both cultivars after chilling stress, which is consistent with previous reports on PSI photoinhibition-inducible stress (Supplementary Fig. S2) (Bernhard Teicher *et al*., 2000; Kılıç *et al*., 2023). This suggested that chilling stress did not inhibit the transcriptional response of Fe-S oxidation pathways in either cultivar.

As photorespiration is a major electron sink in C_3_ plants, photorespiration activity was measured using an oxygen electrode (Supplementary Fig. S3) (Sejima *et al*., 2016; Hanawa *et al*., 2017). However, no differences in photorespiration activity were observed between cultivars under control conditions, and chilling stress significantly reduced photorespiration activity in both cultivars.

As PGR5/PGRL1 is involved in the oxidation of Fe-S, we investigated cultivar differences in PGR5/PGRL1 function by infiltrating cucumber leaves with antimycin A (AA), an inhibitor of PGR5/PGRL1 function and examining P700 redox states before and after chilling stress (Supplementary Fig. S4). Before chilling stress, the AA treatments similarly inhibited the induction of Y(ND) in both cultivars. This trend remained unchanged even after chilling stress, suggesting no cultivar differences in PGR5/PGRL1 function under chilling stress.

The chloroplast NADH dehydrogenase-like complex (NDH) receives electrons from Fd and reduces PQ (Shikanai *et al*., 1998), promoting an oxidation of PSI. NDH activity is characteristically assessed by a transient increase in Chl fluorescence after the AL is turned off (Shikanai *et al*., 1998; Endo *et al*., 1999; Sun *et al*., 2017; Wu *et al*., 2019; Wang *et al*., 2020; He *et al*., 2024; Zhang Y. *et al*., 2024; Ishikawa *et al*., 2016). Under control conditions, interestingly, the chilling-sensitive cultivar HG exhibited lower NDH activity than the chilling-tolerant cultivar HM (Fig. 5A). Furthermore, after chilling stress, NDH activity was no longer detectable in HG (Fig. 5B). To identify the causes of the loss of NDH activity during chilling stress in the chilling-sensitive cultivar HG, thylakoid membrane protein complexes were separated by Blue Native PAGE (Fig. 5C), followed by 2D-SDS-PAGE and immunoblotting with specific antibodies against NDH subunits PnsL1, NdhT, and NdhH (Fig. 5D). The properly assembled NDH complexes were detected as the PSI–NDH supercomplex at the band 1 position (red line). Before chilling stress, NDH signals were mainly detected at the band 1 position in both cultivars. However, after chilling stress, the PnsL1 signal was widely detected at lower molecular weight positions only in the chilling-sensitive cultivar HG (Fig. 5D, red arrow). A similar tendency was observed for NdhT and NdhH, with signals detected at a lower molecular weight only in HG after chilling stress (Fig. 5D). These results demonstrated the partial disassembly and destabilization of the PSI-NDH supercomplex (Opatíková and Kouřil, 2024) by chilling stress in the chilling-sensitive cultivar HG. In contrast, HM maintained the stability of the PSI–NDH supercomplex and retained NDH activity under chilling stress (Fig. 5B, D). In addition, the production of O_2_^•–^ was significantly higher in HG than in HM during the early stages of chilling stress (Fig. 5E). These findings suggested that the chilling-sensitive cultivar HG has lower NDH activity than that in HM even under normal conditions, which makes it susceptible to PSI over-reduction during the initial phase of chilling stress. This leads to greater ROS generation and destabilization of the PSI-NDH supercomplex, ultimately leading to a complete loss of NDH activity and PSI photoinhibition.

**Figure 5.**
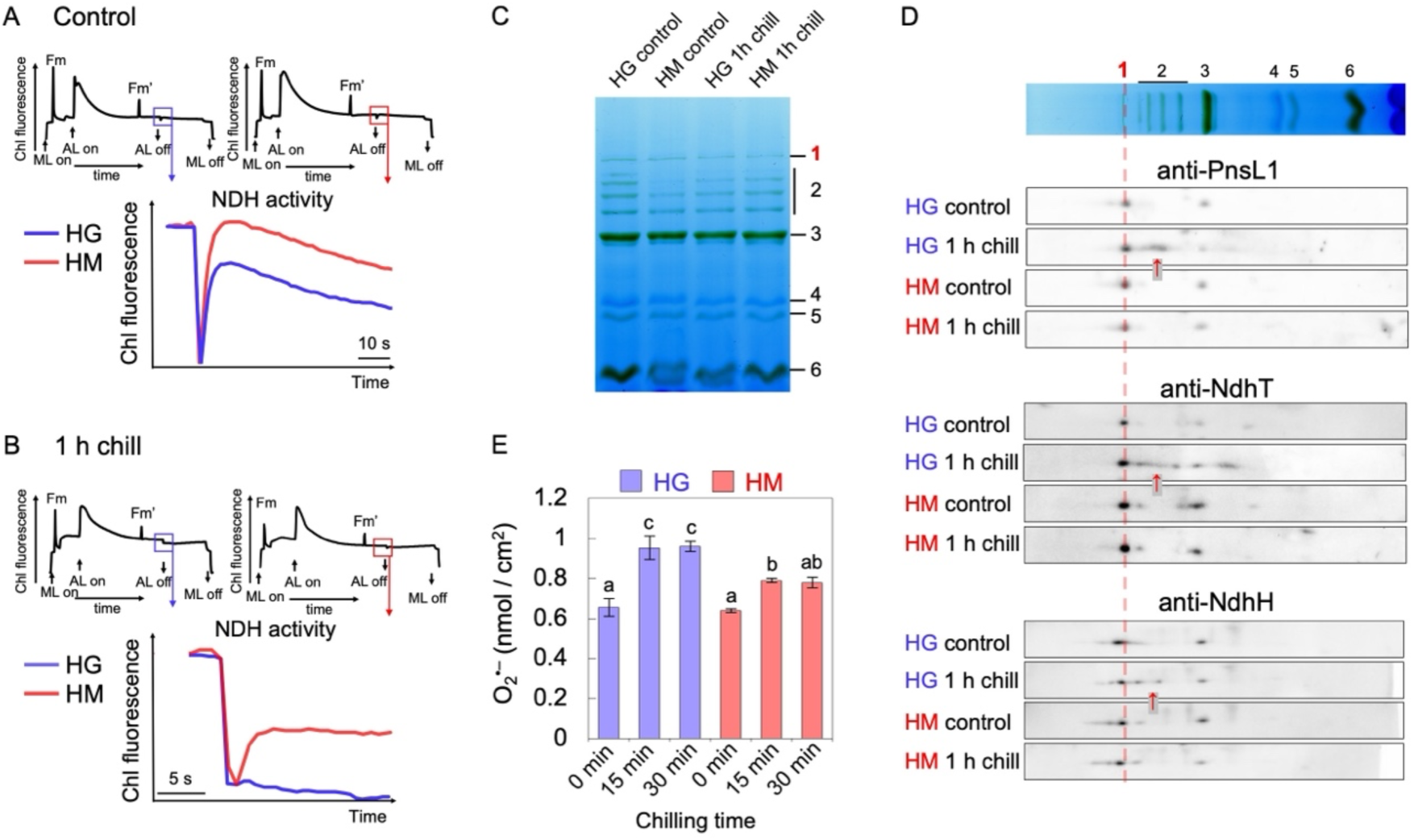
Destabilization of the PSI–NDH supercomplex and ROS generation during chilling stress. (A) Two- to three-week-old cucumbers were treated at 4 °C for 1 h, and NDH activity was analyzed by measuring the transient increase in Chl fluorescence after AL was turned off. Leaves (A) before and (B) after 1 h of chilling stress were dark-adapted for at least 20 min before measurements, followed by exposure to AL for 4–5 min. The subsequent transient increase in Chl fluorescence was monitored in the dark (boxed area). The data shown are typical of the three biological replicates. (C, D) Analysis of the PSI–NDH supercomplex stability before and after 1 h of chilling stress. (C) Thylakoid membrane complexes were separated by BN-PAGE. The isolated thylakoids were solubilized with 1.0% (w/v) n-dodecyl-β-D-maltoside, and the proteins corresponding to 5 µg Chl were separated on a 4–12% gradient gel. (D) Proteins separated by BN-PAGE were analyzed by 2D-SDS-PAGE. NDH subunits (PnsL1, NdhT, and NdhH) were detected using specific antibodies. The red dotted line indicates the PSI–NDH supercomplex band position in BN-PAGE. The BN-PAGE images shown as an example above the immunoblotting data are those obtained with HG control in (C). The red arrows represent the signals of the NDH subunit that have been destabilized by the chilling stress. 1: PSI–NDH supercomplex; 2: PSII–LHCII supercomplex; 3: PSI–LHCI supercomplex, PSII core dimer, and ATPase; 4: PSII core monomer and Cyt *b*_6_*f*; 5: LHCII assembly; and 6: LHCII trimer. (E) Fresh leaves of 2–3-week-old cucumbers were harvested before and immediately after 15 min or 30 min of chilling stress. O_2_^•–^ content was detected as described in Materials and Methods. Values are the mean ± SE, n = 3–4, biological replicates. Different letters indicate statistically significant differences (*p*<0.05, Tukey–Kramer’s multiple comparison tests after one-way ANOVA).

### Impact of differences in NDH activity on PSI

To further elucidate the contribution of NDH to cultivar differences in chilling tolerance, we examined their photosynthetic characteristics under two key conditions that require NDH activity: sudden light illumination and fluctuating light conditions (Shikanai, 2016). Under sudden light illumination (5 min of high light exposure following long dark adaptation), the chilling-sensitive cultivar HG showed delayed P700 oxidation (decreased Y(ND)) and increased Y(NA) without significant changes in PSII parameters (Fig. 6A and Supplementary Fig. S5A). These results closely resembled the general characteristics observed in the NDH-deficient mutant under sudden light illumination (Nikkanen *et al*., 2018; Storti *et al*., 2020; Rodriguez-Heredia *et al*., 2022; Zhou *et al*., 2023). Indeed, the *A. thaliana* NDH-deficient mutant *pnsl1* (Ishihara *et al*., 2007) showed delayed Y(ND) induction and higher Y(NA) than WT plants without significant changes in PSII parameters (Fig. 6B, Supplementary Fig. S5B). Similarly, under fluctuating light conditions, the chilling-sensitive cultivar HG exhibited lower Y(ND) and higher Y(NA) under strong light than did HM, both before and after chilling stress (Fig. 7). This trend is also characteristic of NDH-deficient mutants under fluctuating light conditions (Shikanai, 2016), further supporting our finding that NDH activity was lower in HG than in HM before and during chilling stress (Fig. 5).

**Figure 6.**
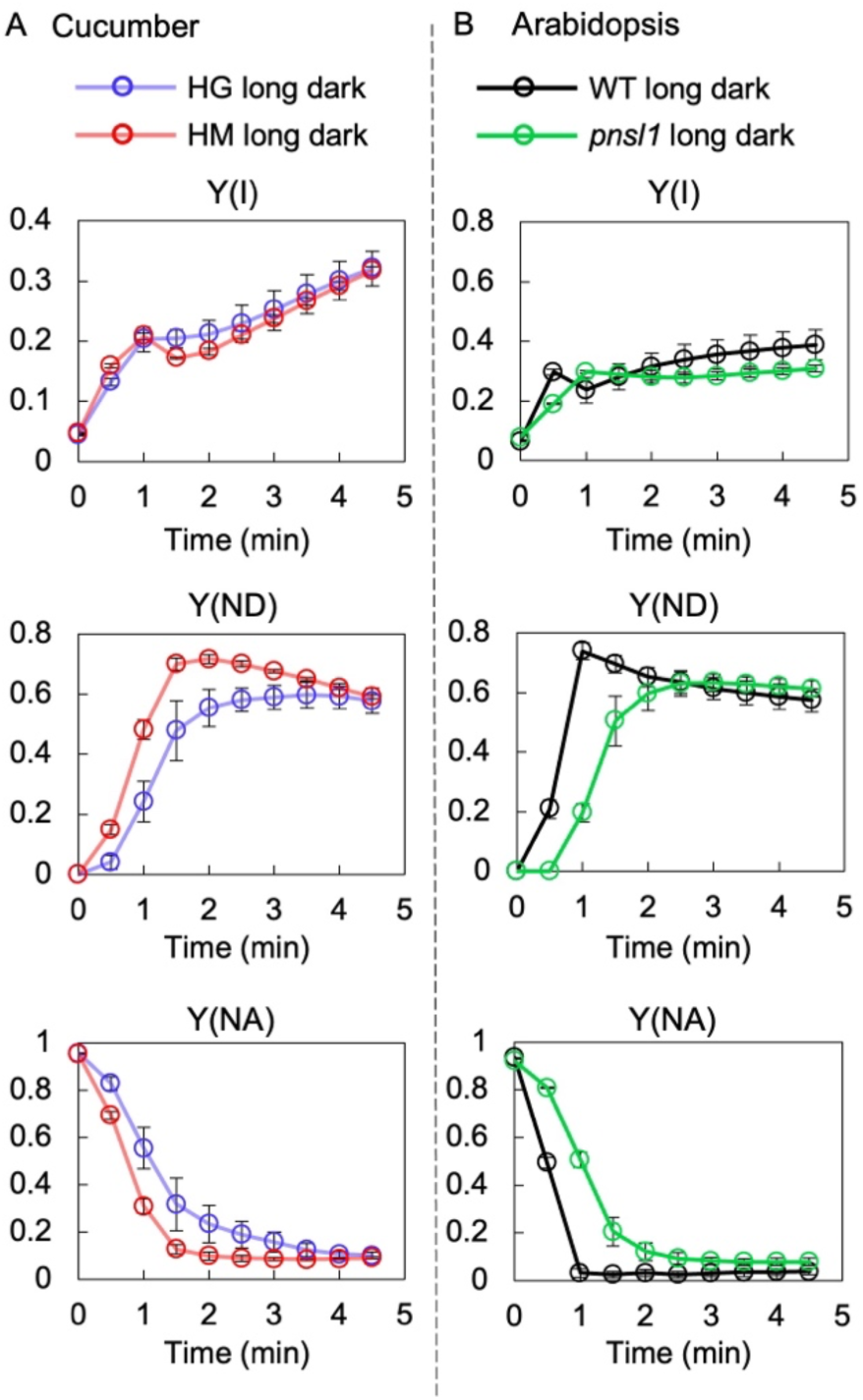
P700 oxidation ability in cucumber and NDH-deficient *Arabidopsis* under sudden light. (A) Chilling-sensitive cv. ‘HG’ and chilling-tolerant cv. ‘HM’ (non-chilled plants) were dark-adapted for 10 h at 23 °C, and the induction phase (dark-to-light transitions) of Chl absorption was monitored under AL (416 µmol photons m^−2^ s^−1^) for 5 min at 23 °C using Dual-PAM 100. (B) *A. thaliana* WT (Col-0) and NDH-deficient mutant *pnsl1* were dark-adapted for 30 min, and the induction phase (dark-to-light transitions) of Chl absorption was monitored under AL (325 µmol photons m^−2^ s^−1^) for 5 min at 23 °C using Dual-PAM 100. Values are the mean ± SE, n = 3–4, biological replicates.

**Figure 7.**
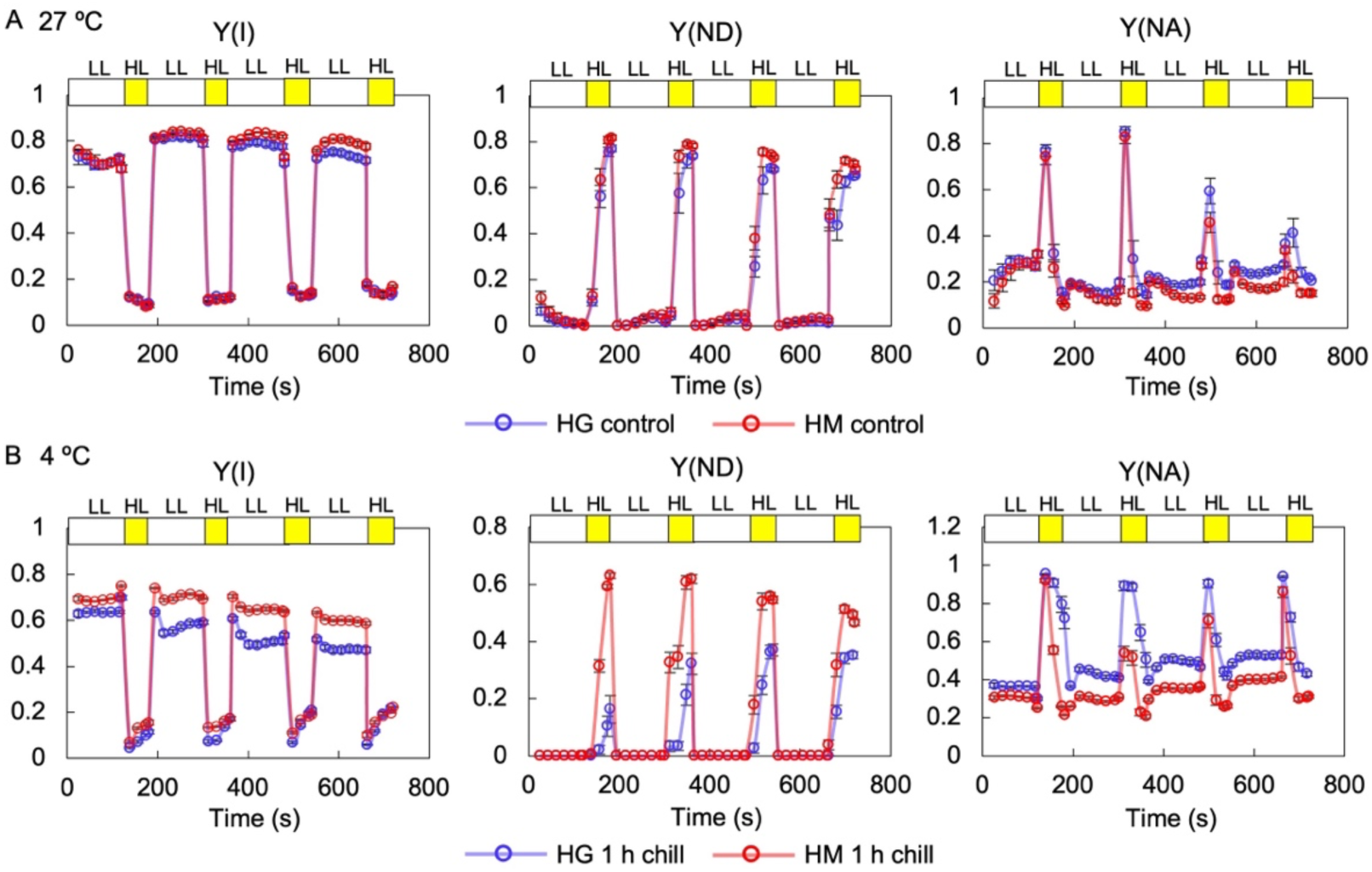
Effects of fluctuating light on PSI in cucumber. (A) Non-chilled 2–3-week-old cucumbers (chilling-sensitive cv. ‘HG’ and chilling-tolerant cv. ‘HM’) were exposed to fluctuating light conditions repeating low light (LL; 15 µmol photons m^−2^ s^−1^ for 2 min, indicated by open bars) and high light (HL; 1,400 µmol photons m^−2^ s^−1^ for 1 min, indicated by yellow bars) at 23 °C. Measurements were taken after at least 20 min of dark adaptation. (B) Cucumbers were exposed to 1 h of chilling stress, and subsequently, they were exposed to the same fluctuating light conditions as mentioned in (A). Values are the mean ± SE, n = 3–4, biological replicates.

Finally, as PSI was more over-reduced under CO_2_-limited conditions in the chilling-sensitive cultivar HG than in the chilling-tolerant cultivar HM (Fig. 4), we confirmed the role of NDH as an electron sink other than the CBB cycle downstream of PSI by monitoring the P700 redox states in the *A. thaliana* NDH-deficient mutant *pnsl1* under low CO_2_ conditions. *pnsl1* exhibited lower Y(ND) and significantly higher Y(NA) than WT under CO_2_-limited conditions, indicating severe PSI over-reduction (Fig. 8). No differences were found in PSII parameters, confirming that electron influx into PSI was similar between the WT and *pnsl1* (Fig. 8). These differences in over-reduction of PSI between WT and *pnsl1* were similar to the cultivar differences in cucumber (Fig. 4). These results demonstrated that NDH functions as an electron sink downstream of PSI under CO_2_-limited conditions and emphasized its importance in protecting PSI from photoinhibition during chilling stress when the CBB cycle is inhibited (Kingston-Smith *et al*., 1997; Zhang Q. *et al*., 2024).

**Figure 8.**
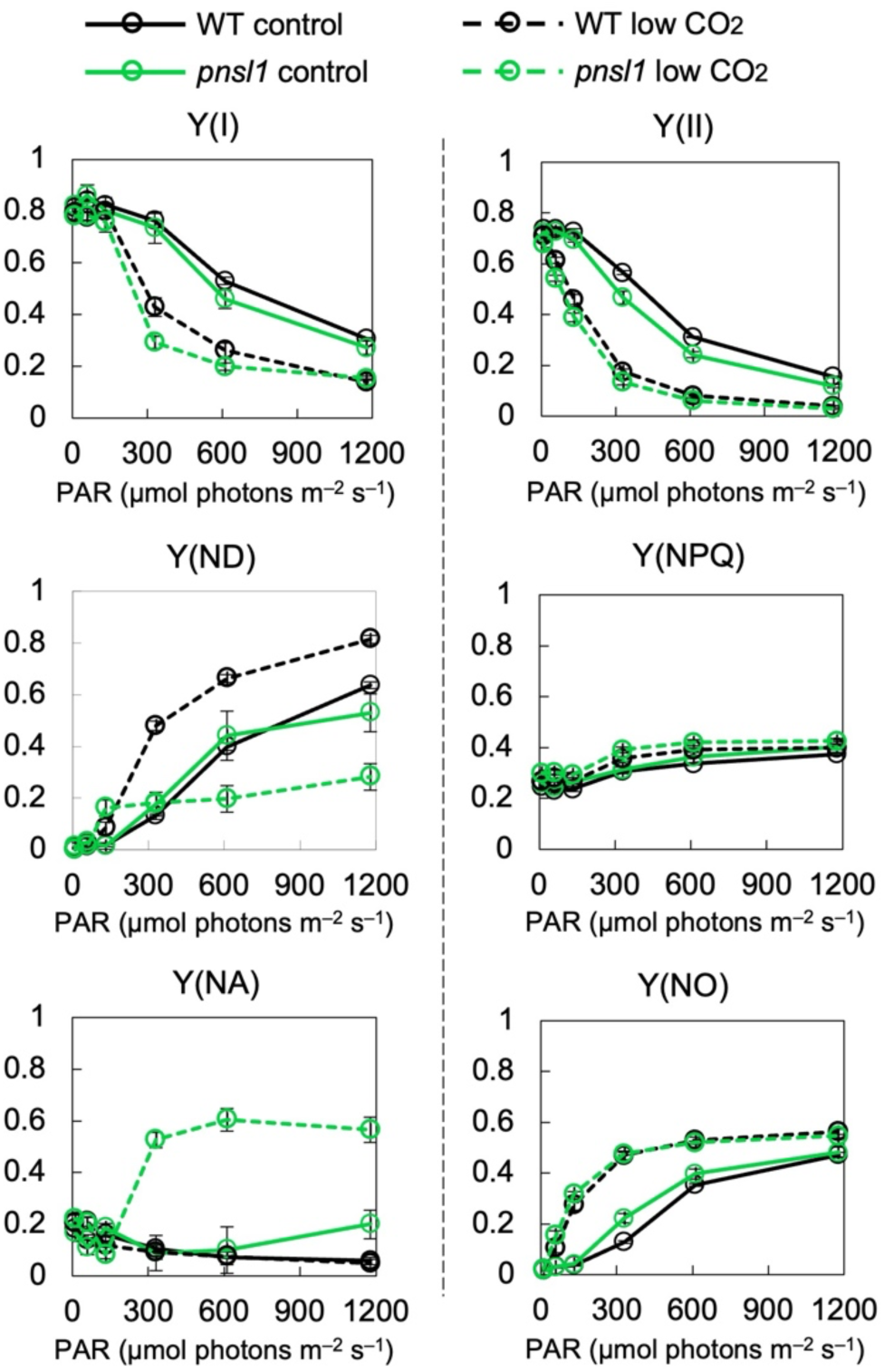
Impact of CO_2_ limitation on P700 oxidation in NDH-deficient *Arabidopsis*. Chl fluorescence and P700 redox states were measured under various light intensities in ambient (control) or low CO_2_ (<30 ppm) conditions at 23 °C using Dual-PAM 100. The intact leaves of *A. thaliana* WT (Col-0) and the NDH-deficient mutant *pnsl1* were analyzed. Values are the mean ± SE, n = 3–4, biological replicates.

Altogether, in the chilling-tolerant cucumber, NDH functions effectively as an electron sink, maintaining P700 and Fe-S oxidized, thereby suppressing ROS generation and mitigating the damage of chloroplast ATPase and PSI photoinhibition (Fig. 9A). In contrast, insufficient NDH activity led to over-reduction of the PSI acceptor side up to Fd, exacerbating PSI photoinhibition induced by chilling stress (Fig. 9B).

**Figure 9.**
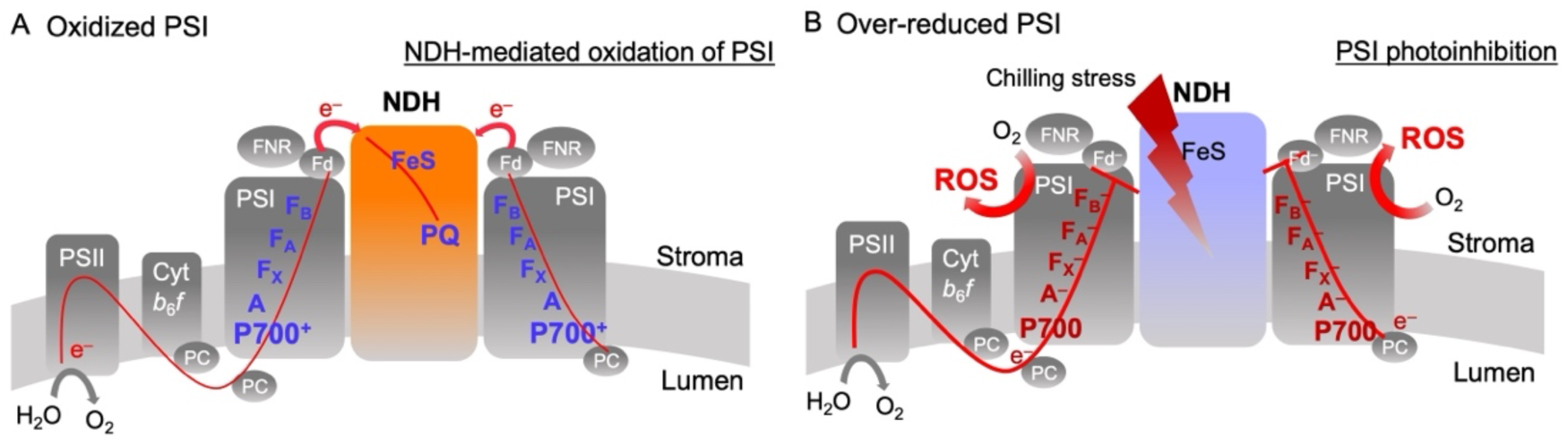
Importance of NDH in preventing PSI photoinhibition. (A) PSI electron transfer pathway follows P700→A_0_→A_1_→F_X_→F_A_/F_B_→Fd, where A_0_ is the primary acceptor Chl *a*, A_1_ is the secondary acceptor phylloquinone, and F_X_, F_A_, and F_B_ are Fe-S clusters. Under suppressed CBB cycle conditions, such as chilling stress, NDH protects PSI from its over-reduction by accepting electrons from reduced Fd. A reduced Fd molecule binds to the subcomplex E of NDH in the stroma, and three Fe-S clusters within NDH facilitate electron transfer. One PQ molecule within the NDH complex can accept two electrons, enabling another reduced Fd to bind immediately. This electron acceptance by NDH keeps A_0_, A_1_, F_X_, and F_A_/F_B_ in an oxidative state, thereby suppressing ROS generation. (B) In chilling-sensitive cucumbers, the PSI–NDH supercomplex is destabilized by chilling stress and cannot function as an electron acceptor for Fd. Consequently, PSI becomes over-reduced, and excess electrons accumulate at PSI, leading to ROS generation and ultimately to PSI photoinhibition.

## Discussion

In this study, we examined two cucumber cultivars that exhibited different levels of chilling stress tolerance and PSI photoinhibition (Figure 1) to elucidate the cause of PSI photoinhibition. Under chilling stress, cucumbers undergo thylakoid membrane uncoupling, leading to incomplete photosynthetic control (Fig. 2) (Kono *et al*., 2022). Notably, the P700 and Fe-S clusters were over-reduced only in the chilling-sensitive cultivar (Fig. 1C and 3A) because of the suppression of electron efflux from Fd (Fig. 3B). Furthermore, in the chilling-sensitive cultivars, PSI was over-reduced not only under chilling stress, but also during CO_2_-limited conditions (Fig. 4), suggesting the importance of AEF pathways downstream of Fd. Among the various AEF pathways, NDH activity differed significantly between the cultivars (Fig. 5A, B) NDH activity was lower in the chilling-sensitive cultivar than in the chilling-tolerant cultivar and was completely lost under chilling stress because of the destabilization of PSI–NDH supercomplexes (Fig. 5D). The over-reduction of PSI during dark–light transitions (Fig. 6) and under fluctuating light (Fig. 7) further supports the importance of NDH in PSI photoinhibition under chilling stress. Our findings demonstrated that impaired NDH function under chilling stress leads to Fe-S over-reduction (Fig. 3), ultimately resulting in PSI photoinhibition. This study reveals the crucial role of NDH in chilling stress tolerance and the prevention of PSI photoinhibition in land plants (Fig. 9).

In angiosperms, the NDH complex consists of at least 29 subunits organized into five subcomplexes (Ifuku *et al*., 2011): subcomplexes M (SubM) (NdhA–NdhG), SubA (NdhH–NdhO), SubE (NdhS–NdhV), SubB (PnsB1–PnsB5), and the luminal subcomplex (SubL) (PnsL1–PnsL5). SubL is unique to angiosperms, and SubE forms the Fd-binding site (Schuller *et al*., 2019). The PSI–NDH supercomplex consists of a single NDH sandwiched between two PSI units (Kouřil *et al*., 2014; Yadav *et al*., 2017; Shen *et al*., 2022). Its assembly largely depends on LHCA proteins, in particular Lhca5 and Lhca6 (Otani *et al*., 2017, 2018; Kato *et al*., 2018; Shen *et al*., 2022; Su *et al*., 2022; Opatíková and Kouřil, 2024; Introini *et al*., 2025). Recent advances in cryo-electron microscopy (cryo-EM) have expanded our understanding of NDH structure and biochemical functions. Once dissociated from PSI, Fd binds to SubE and transfers electrons through three Fe-S clusters embedded within the NDH, ultimately reducing the PQ molecule inside the NDH complex (Strand *et al*., 2017; Schuller *et al*., 2019; Pan *et al*., 2020; Richardson *et al*., 2021; Su *et al*., 2022; Introini *et al*., 2025). The interaction between Fd and NDH was as robust as that between PSI and Fd. This strong interaction and the short distance between PSI and NDH suggest that Fd efficiently transfers electrons from PSI to NDH within the supercomplex (Su *et al*., 2022). In addition, the structural changes in NDH with pH (Pan *et al*., 2020) and the activation of NDH by H_2_O_2_, heat, and the thioredoxin system (Casano *et al*., 2001; Lascano *et al*., 2003; Nikkanen *et al*., 2018; Zhang *et al*., 2023) strongly suggest a potential physiological role for NDH. However, NDH-deficient mutants typically exhibit phenotypes similar to those of wild-type in C_3_ plants, and their physiological significance in response to environmental stress is largely unclear (Yamori *et al*., 2011; Urzinger *et al*., 2025).

This study identified two novel physiological roles for NDH. First, under CO₂-limited conditions, NDH functions as an electron sink, preventing PSI over-reduction, as evidenced by the NDH-deficient *A. thaliana* mutant exhibiting enhanced PSI over-reduction under CBB cycle-limited conditions (Fig. 8). Second, under chilling stress, NDH was essential for alleviating Fe-S over-reduction and suppressing PSI photoinhibition (Fig. 3 and 5). Both of these NDH functions became evident when the CBB cycle was restricted. Under these conditions, NDH accepts electrons from Fd and reduces plastoquinone (PQ) via three Fe-S clusters within the NDH complex, thereby contributing to the redistribution of reducing power downstream of PSI. Furthermore, the increase in reduced PQ via NDH inhibited electron transfer at Cyt *b*_6_*f* independently of ΔpH, leading to P700 oxidation. This mechanism is consistent with the concept of the reduction-induced suppression of electron flow (RISE) (Shaku *et al*., 2016).

The two cucumber cultivars used in this study exhibited different responses to sudden light and fluctuating light, even under normal temperatures (Figs 6 and 7), and their NDH activities differed under ambient conditions (Fig. 5A, B). We compared the amino acid sequences of all nuclear-encoded NDH subunits between the chilling-sensitive cultivar HG and the chilling-tolerant cultivar HM. However, the sequences were highly conserved between cultivars, except for the residue Ala162 of NdhT (Ala162 to Val162) (Supplementary Fig. S6A). Although the properties of Ala and Val are relatively similar, this residue is located near the interaction domain with NdhJ (Supplementary Fig. S6B). Although studies on the impact of specific amino acid residues of NDH on its activity remain limited, recent research on a single amino acid substitution in NdhF has demonstrated that the mutation of “Asp635” to “Ala635” alters interactions between NdhF and NdhD, leading to decreased NDH activity and changes in complex stability in cyanobacteria (Zheng *et al*., 2025). However, further research is needed to elucidate the regulatory mechanisms of NDH activity through changes from Ala162 to Val162 in NdhT and their evolutionary significance.

In the chilling-sensitive cultivar with lower NDH activity under normal conditions, ROS generation was more pronounced under stress conditions (Fig. 5E). This aligns with previous reports showing that NDH-deficient tobacco and cyanobacteria mutants exhibit increased ROS production under stress (Zhang *et al*., 2023; Zheng *et al*., 2025). In addition, exposure to chilling stress led to the destabilization of supercomplexes and a complete loss of NDH activity in this cultivar (Fig. 5). We propose that ROS generated under stress conditions attack the PSI–LHCI–NDH supercomplex, ultimately leading to its destabilization (Fig. 5D), because the interaction between PSI–LHCI and NDH is relatively unstable (Su *et al*., 2022; Opatíková and Kouřil, 2024) and because PSI photoinhibition is associated with damages to LHCAs (Alboresi *et al*., 2009; Takagi *et al*., 2022). Interestingly, P700 oxidation by far-red (FR) light does not occur after chilling-induced PSI photoinhibition (Havaux and Davaud, 1994; Sonoike, 1999). This strongly implies damages to LHCAs and NDH under light-chilling stress, as low-energy Chls in LHCAs are essential for FR light absorption, and NDH contributes to P700 oxidation under FR light (Kono *et al*., 2017). The stability of LHCAs under chilling stress, particularly Lhca5 and Lhca6, needs to be elucidated in future studies. In addition, Fe-S clusters within NDH are highly reactive to ROS (Petronek *et al*., 2021), potentially triggering NDH degradation under photo-oxidative stress.

The role of NDH in environmental stress tolerance has been overlooked because phenotypic changes caused by NDH deficiency appear only under fluctuating light conditions in the model plant *A. thaliana*. This underestimation was probably because of the varying importance of NDH in various plant species and ecotypes adapted to different environments. In *A. thaliana*, chilling stress rapidly decreases Fv/Fm and leads to ΔpH-driven PSII inactivation, followed by PSI oxidation (Takeuchi *et al*., 2025). As a result, the Fe-S clusters do not reach the reduction level, requiring NDH-mediated electron dissipation. In contrast, cucumbers did not immediately experience PSII inactivation under chilling stress. The decrease in Fv/Fm occurred later than that in Pm (Fig. 1B) (Takeuchi *et al*., 2025). Consequently, a mechanism that alleviates PSI over-reduction on the PSI acceptor side—NDH activity—is essential. In chilling-tolerant cucumbers, NDH activity was constitutively high (Fig. 5–7), enabling the effective dissipation of PSI over-reduction under chilling stress (Fig. 1D and 3). In addition, 6 units of PSI assembled for 1 unit of NDH complex have been observed in *A. thaliana* (PSI : NDH = 6:1) (Yadav *et al*., 2017). The amount of NDH around PSI may be differently regulated in various plant species and environments. Further research is required to elucidate the differences in the importance of NDH in various plant species.

For many plant species, the natural situation that frequently results in PSI photoinhibition is fluctuating light because sunlight during the day is highly variable. Fluctuating light stress can easily induce PSI photoinhibition even in WT plants (Kono and Terashima, 2016; Yamori *et al*., 2016; Kono *et al*., 2025). However, PSI photoinhibition is less severe in the field. FR light is a key factor in preventing PSI photoinhibition in nature. It activates NDH and protects PSI from fluctuating light-induced photoinhibition (Kono *et al*., 2017). In natural environments, where FR light is much more abundant, FR light enhances NDH activity and suppresses PSI photoinhibition (Kono *et al*., 2022). Although NDH mutants show minimal phenotypic differences under controlled laboratory conditions, NDH must play an important role in suppressing PSI photoinhibition and environmental stress tolerance in natural environments.

## Acknowledgments

We sincerely thank Dr. Masaru Kono (Astrobiology Center, Japan) and Dr. Daisuke Takagi (Setsunan Univ., Japan) for their valuable discussions and insightful suggestions. This work was supported in part by CREST, Japan Science and Technology Agency, grant number (JPMJCR15O3, JPMJCR17O2 to K.I.) and by Grant-in-Aid for Japan Society for the Promotion of Science Fellows, grant number (JP23KJ1357 to K.T.).

## Author contributions

Ko Takeuchi, Shintaro Harimoto, Yufen Che, and Kentaro Ifuku designed the research. Ko Takeuchi, Shintaro Harimoto, Yufen Che, and Minoru Kumazawa performed the experiments. Ko Takeuchi, Kentaro Ifuku, Chikahiro Miyake, Shintaro Harimoto, Yufen Che, Hayato Satoh, and Shu Maekawa analyzed the data. Chikahiro Miyake provided the Dual-KLAS-NIR. Ko Takeuchi and Kentaro Ifuku wrote the manuscript.

## Data availability

The data underlying this article are available in the article and its online supplementary material.

## Supporting Information

Additional Supporting Information may be found online in the Supporting Information section at the end of the article.

Fig. S1 Separation of thylakoid membrane complexes by CN-PAGE.

Fig. S2 Gene expression analysis of pathways involved in Fd oxidation.

Fig. S3 Photorespiration activity before and after chilling stress.

Fig. S4 Effects of antimycin A (AA) on P700 oxidation before and after chilling stress.

Fig. S5 Effects of sudden light on PSII parameters.

Fig. S6 Comparison of the amino acid sequence of NdhT between cucumber cultivars.

