## Supporting Information for "The protective role of chloroplast NADH dehydrogenase-like complex (NDH) against PSI photoinhibition under chilling stress"

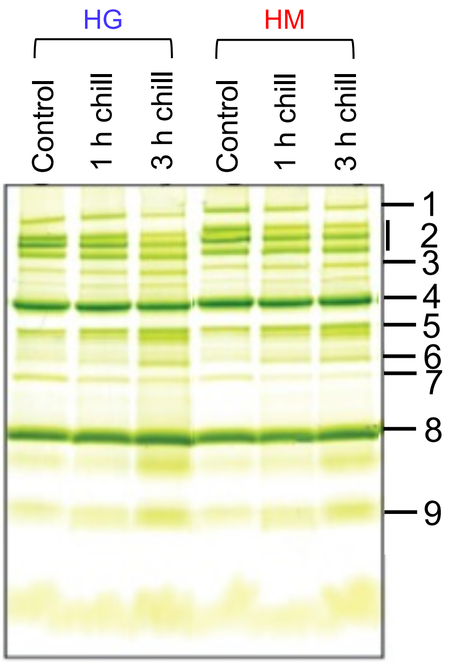


**Supplementary Figure S1. Separation of thylakoid membrane complexes by CN-PAGE.** Two- to three-week-old cucumbers were treated at 4 ºC for 1 h or 3 h. The isolated thylakoids (10 µg Chl) were solubilized using 1% β-DM and separated by CN-PAGE. 1: PSI–NDH supercomplex; 2: PSII–LHCII supercomplex; 3: PSI + LHCI; 4: PSI–LHCI, PSII core dimer, and ATPase; 5: PSI core monomer; 6: PSII core monomer and Cyt $b_6f$; 7: LHCII assembly; 8: LHCII trimer; and 9: LHCII monomer.

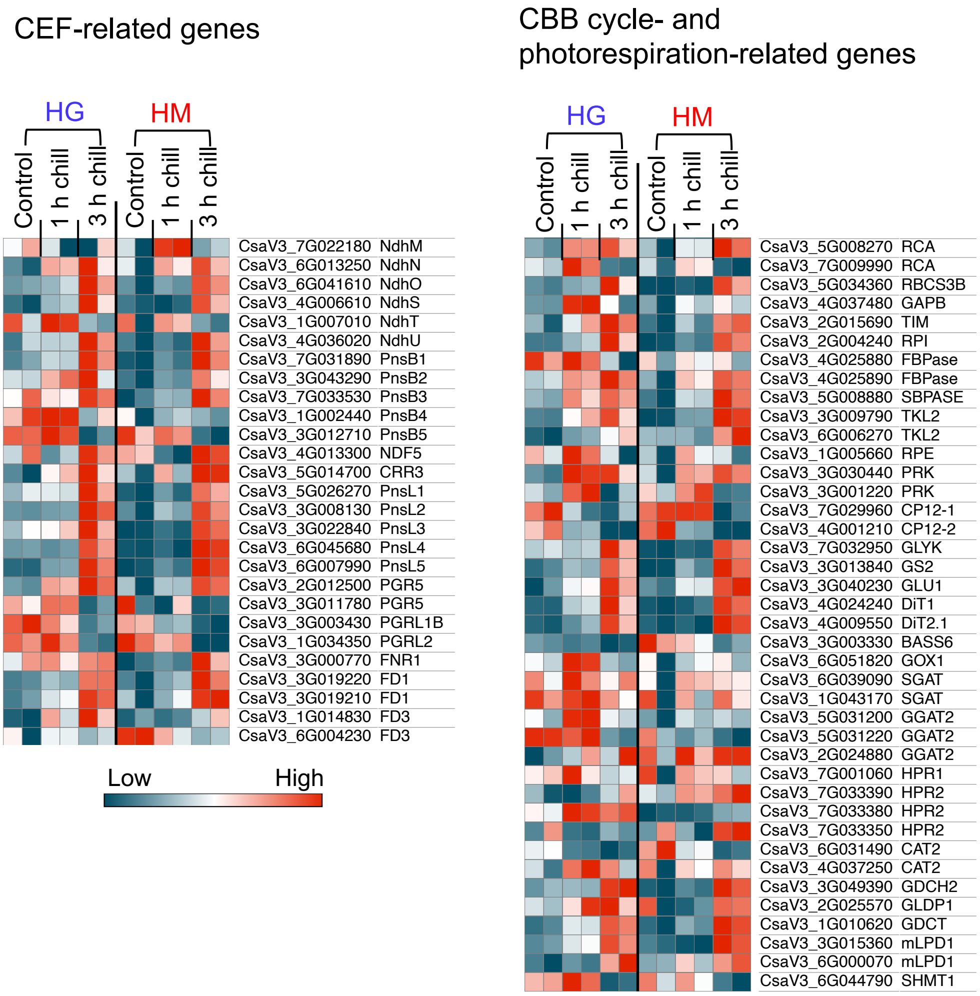


**Supplementary Figure S2. Gene expression analysis of pathways involved in Fd oxidation.** All nuclear-coded CEF-, CBB cycle-, and photorespiration-related genes were analyzed by RNA-seq. The gene list was obtained from a published report (Kılıç *et al.*, 2023), and a heatmap was generated using Morpheus. RNA was extracted from 2–3-week-old cucumber leaves before and after 1 h or 3 h of chilling stress. n = 2, biological replicates.

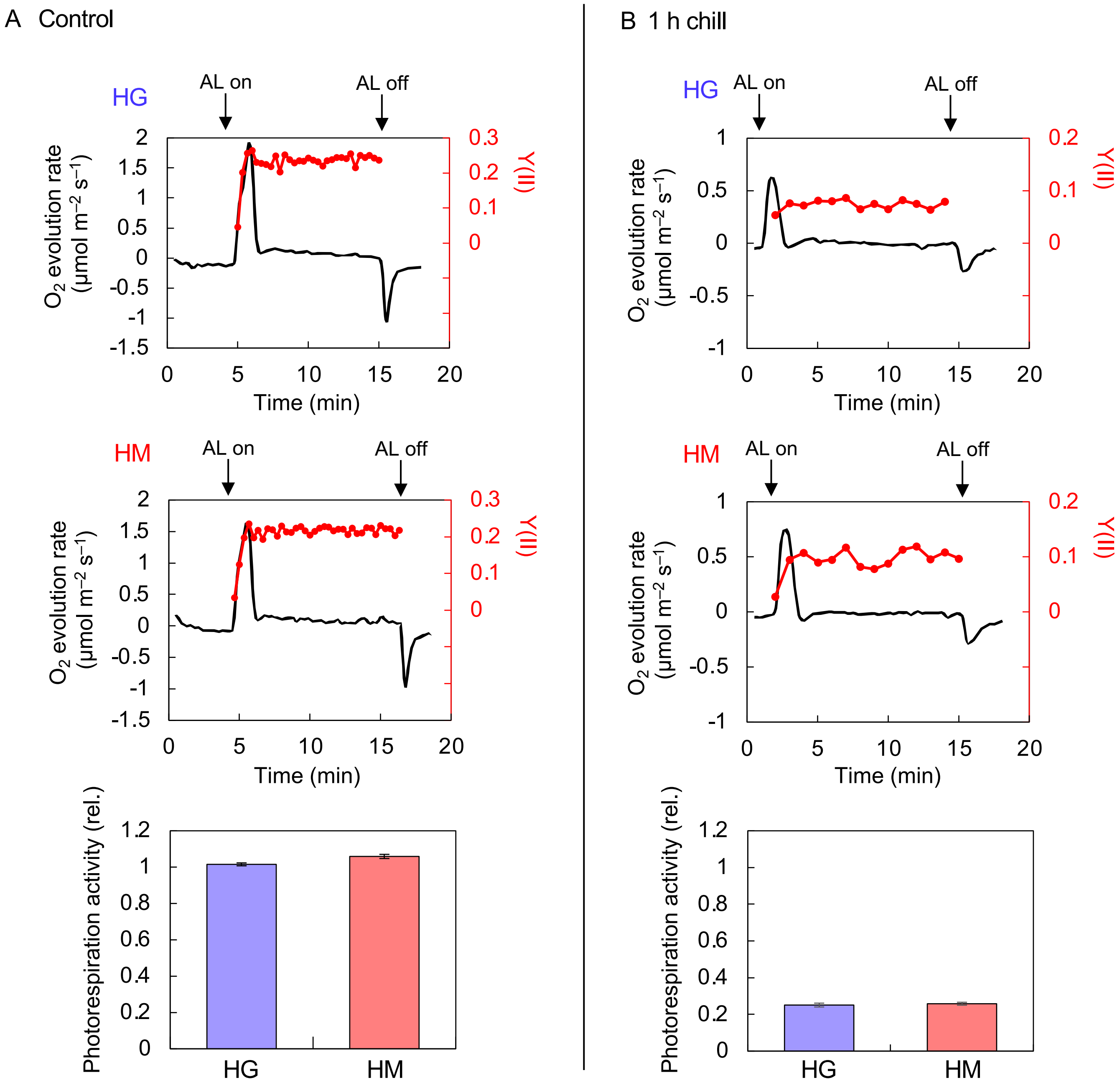


**Supplementary Figure S3. Photorespiration activity before and after chilling stress.** Photorespiration activity was measured as described in Materials and Methods. $O_2$ exchange was monitored simultaneously with Chl fluorescence. A leaf disc was placed in an $O_2$-electrode chamber, and Chl fluorescence was monitored using Junior-PAM. The post-illumination transient $O_2$-uptake rate reflected the activities of RuBP oxygenase. (A) Before chilling stress (control) and (B) After chilling stress for 1 h. $O_2$ evolution rate and Y(II) kinetics show a typical series of three biological replicates. Values are the mean ± SE, n = 3, biological replicates.

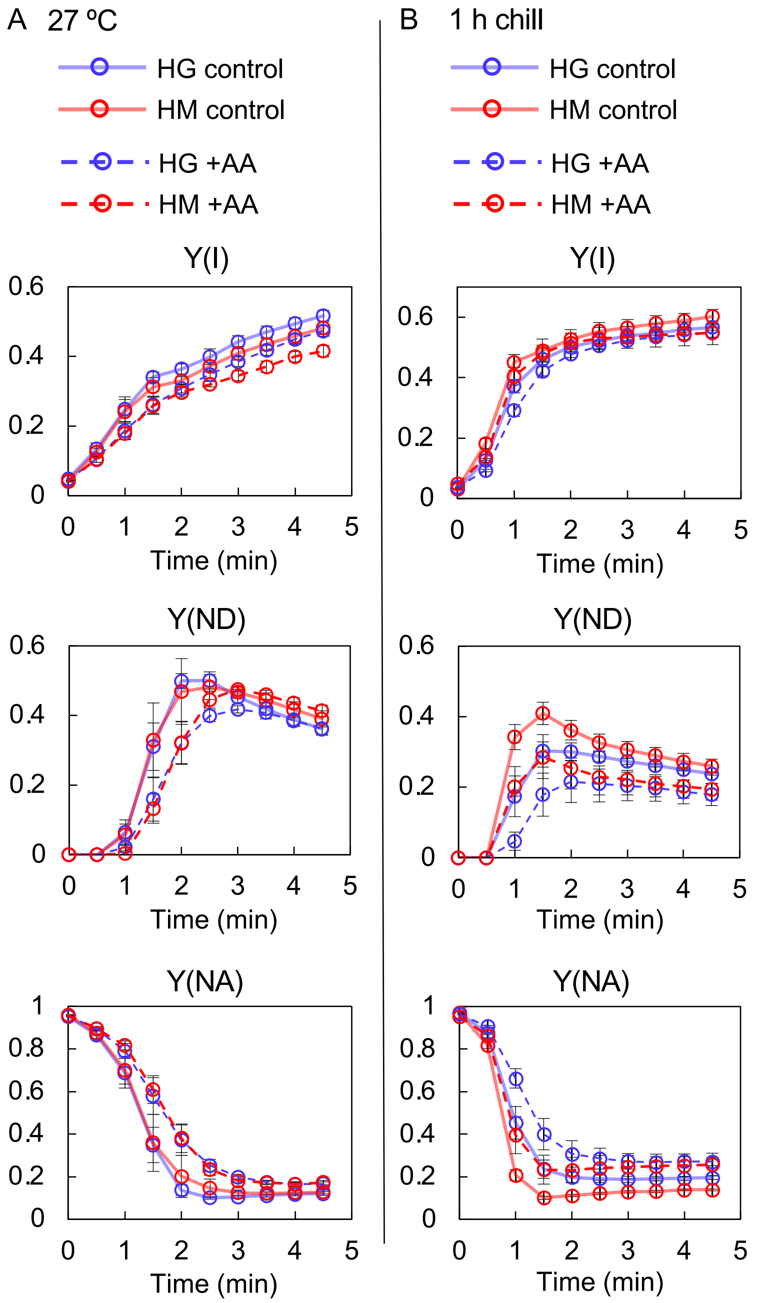


**Supplementary Figure S4. Effects of antimycin A (AA) on P700 oxidation before and after chilling stress.** Chl fluorescence and P700 redox states were analyzed in 2–3-week-old cucumber plants (chilling-sensitive cv. 'HG' and chilling-tolerant cv. 'HM') (A) before and (B) after 1 h of chilling stress. Intact leaves were infiltrated with 20 µM AA in ethanol using desiccator (−0.1 MPa for 2 min). Subsequently, the leaves were immersed in 20 µM AA solution, shaken slowly in the dark for an hour, and then exposed to chilling stress. Chilling-treated plants were dark-adapted for at least 20 min at 4 ºC, whereas control plants were adapted at 23 ºC. Chl fluorescence and absorption under the induction phase (dark-to-light transitions) of photosynthesis were measured using Dual-PAM 100. AL (325 µmol photons $m^{-2}$ $s^{-1}$) was applied under ambient conditions (23 ºC) for 5 min. Values are the mean ± SE, n = 3, biological replicates.

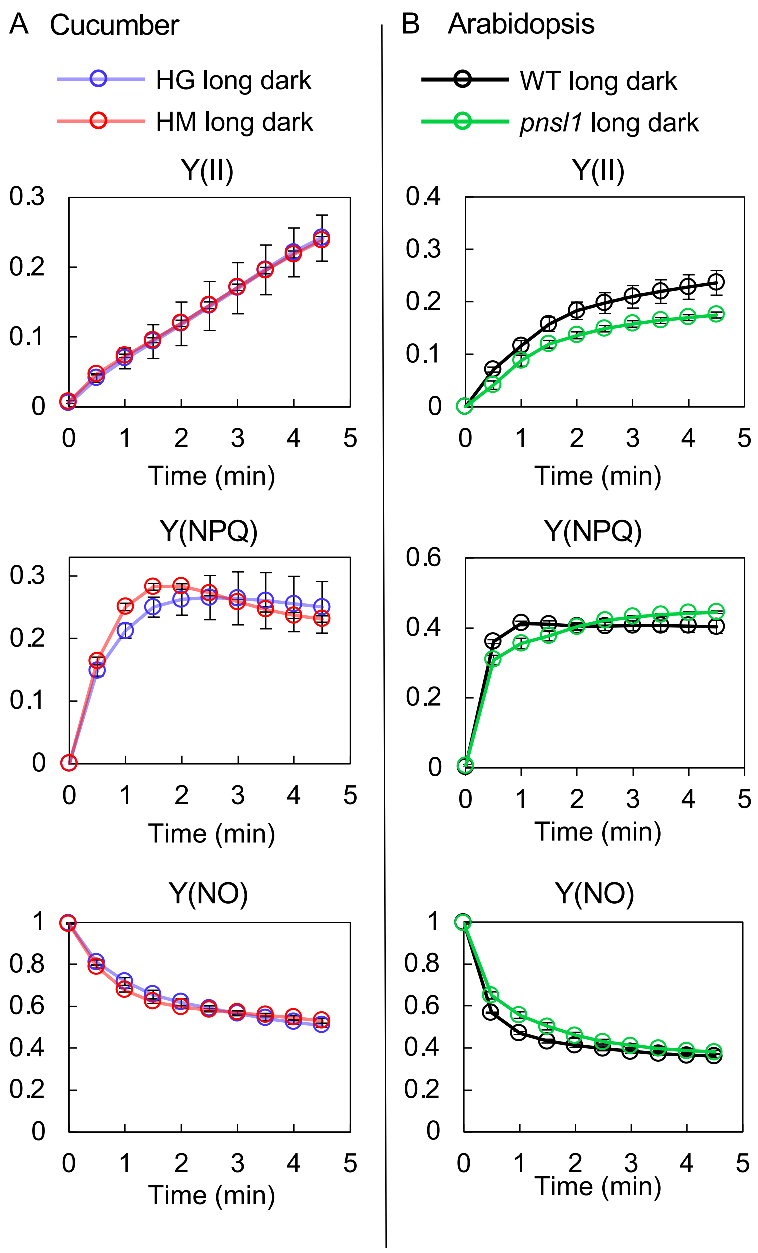


**Supplementary Figure S5. Effects of sudden light on PSII parameters.** The plants and experimental conditions were the same as mentioned in Figure 6. (A) Intact leaves of 2–3-week-old cucumbers (chilling-sensitive cv. ‘HG’ and chilling-tolerant cv. ‘HM’). (B) *A. thaliana* WT (Col-0) and NDH-deficient mutant *pnsl1*. Values are the mean ± SE, n = 3–4, biological replicates.

A

1 50

MASPPFPNSLLFHKTFHSSAPTFRTWRVYAAQAPNNDRRRPPPGVDTRIHWQNEEEG
MASPPFPNSLLFHKTFHSSAPTFRTWRVYAAQAPNNDRRRPPPGVDTRIHWQNEEEG

100

WIGRKKKESNDQNNVSNNMLGPSLADLLNNSSDSHYQFLGVDAEAEVEEIKSAYRRLS
WIGRKKKESNDQNNVSNNMLGPSLADLLNNSSDSHYQFLGVDAEAEVEEIKSAYRRLS

150 163

KEYHPDTTSLPLKVASEKFMKLKQVYEVLSNEESRKFYDWTLAQEEVSRQADKLRMKL
KEYHPDTTSLPLKVASEKFMKLKQVYEVLSNEESRKFYDWTLAQEEASRQADKLRMKL
*

200 234

EDPYEQELQNWVSTPDMVDRLGGRNLELSDQASTALTLDIFIILFAIACIVYVLLFKEPYY
EDPYEQELQNWVSTPDMVDRLGGRNLELSDQASTALTLDIFIILFAIACIVYVLLFKEPYY

B

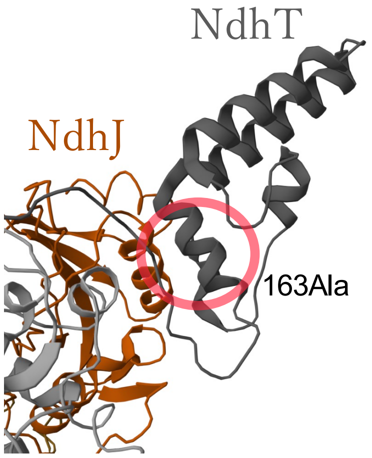


**Supplementary Figure S6. Comparison of the amino acid sequence of NdhT between cucumber cultivars.** The amino acid sequences of the cucumbers (chilling-sensitive cv. 'HG' and chilling-tolerant cv. 'HM') were estimated from RNA-seq data. Of all nuclear-encoded NDH genes, NdhT differed only at the amino acid position 163 between the cultivars, with HG and HM having Ala and Val at this position (A). 163Ala is located near the binding region to NdhJ (B). The structure data was obtained from the PDB (ID: 7WG5).
